# Bayesian Comparison of Explicit and Implicit Causal Inference Strategies in Multisensory Heading Perception

**DOI:** 10.1101/150052

**Authors:** Luigi Acerbi, Kalpana Dokka, Dora E. Angelaki, Wei Ji Ma

**Affiliations:** New York University; Baylor College of Medicine

**Keywords:** Bayesian, model fitting, model comparison, causal inference, multisensory, heading

## Abstract

The precision of multisensory heading perception improves when visual and vestibular cues arising from the same cause, namely motion of the observer through a stationary environment, are integrated. Thus, in order to determine how the cues should be processed, the brain must infer the causal relationship underlying the multisensory cues. In heading perception, however, it is unclear whether observers follow the Bayesian strategy, a simpler non-Bayesian heuristic, or even perform causal inference at all. We developed an efficient and robust computational framework to perform Bayesian model comparison of causal inference strategies, which incorporates a number of alternative assumptions about the observers. With this framework, we investigated whether human observers’ performance in an *explicit* cause attribution and an *implicit* heading discrimination task can be modeled as a causal inference process. In the explicit inference task, all subjects accounted for cue disparity when reporting judgments of common cause, although not necessarily all in a Bayesian fashion. By contrast, but in agreement with previous findings, data from the heading discrimination task only could not rule out that several of the same observers were adopting a forced-fusion strategy, whereby cues are integrated regardless of disparity. Only when we combined evidence from both tasks we were able to rule out forced-fusion in the heading discrimination task. Crucially, findings were robust across a number of variants of models and analyses. Our results demonstrate that our proposed computational framework allows researchers to ask complex questions within a rigorous Bayesian framework that accounts for parameter and model uncertainty.

## Introduction

We constantly interact with people, animals and objects around us. As a consequence, our central nervous system (CNS) receives sensory inputs from multiple modalities (e.g., visual, auditory, vestibular, proprioceptive) that arise from the same or different events in the world. For efficient interaction with the world, the CNS must decide whether the multisensory cues originated from the same cause and should be integrated into a single percept or cues should be interpreted in isolation as they arose from different causes (segregation). Despite the abundance of sensory information, we typically are able to integrate relevant cues while ignoring irrelevant sensory input. It is thus plausible that our brain infers the causal relationship between multisensory cues to determine if and how the cues should be integrated.

Bayesian causal inference – inference of the causal relationship between observed cues, based on the inversion of the statistical model of the task – has been proposed as the decision strategy adopted by the CNS to address the problem of integration vs. segregation of sensory cues (Körding et al., 2007). Such a decision strategy has described human performance in spatial localization (Beierholm, Quartz, & Shams, 2009; Bejjanki, Knill, & Aslin, 2016; Körding et al., 2007; Odegaard & Shams, 2016; Odegaard, Wozny, & Shams, 2015; Rohe & Noppeney, 2015a, 2015b; Sato, Toyoizumi, & Aihara, 2007; Wozny, Beierholm, & Shams, 2010; Wozny & Shams, 2011), orientation judgment van den Berg, Vogel, Josić, and Ma (2012), oddity detection (Hospedales & Vijayakumar, 2009), speech perception (Magnotti, Ma, & Beauchamp, 2013) and time-interval perception (Sawai, Sato, & Aihara, 2012). Over the years, Bayesian models have become more complex as they include more precise descriptions of the sensory noise (de Winkel, Katliar, & Bülthoff, 2015, 2017; Odegaard et al., 2015) and alternative Bayesian decision strategies (Rohe & Noppeney, 2015b; Wozny et al., 2010). However, it is still unknown whether observers fully implement Bayesian causal inference, or merely an approximation that does not take into account the full statistical structure of the task. For example, the Bayes-optimal inference strategy ought to incorporate sensory uncertainty into its decision rule. On the other hand, a suboptimal heuristic decision rule may disregard sensory uncertainty (Ma, 2012; Qamar et al., 2013). Thus, the growing complexity of models and the need to consider alternative hypotheses require an efficient computational framework to address these open questions while avoiding trappings such as overfitting or lack of model identifiability (Acerbi, Ma, & Vijayakumar, 2014).

Visuo-vestibular integration in heading perception presents an ideal case to characterize the details of the causal inference strategy in multisensory perception. While a wealth of published studies have shown that integration of visual and vestibular self-motion cues increases perceptual precision (Butler, Campos, Bülthoff, & Smith, 2011; Butler, Smith, Campos, & Bülthoff, 2010; de Winkel et al., 2013; de Winkel, Weesie, Werkhoven, & Groen, 2010; Dokka, DeAngelis, & Angelaki, 2015; Fetsch, Turner, DeAngelis, & Angelaki, 2009; Gu, Angelaki, & DeAngelis, 2008; Prsa, Gale, & Blanke, 2012), such an integration only makes sense if the two cues arise from the same cause – that is movement of the observer through a stationary visual environment. Despite the putative relevance of causal inference in heading perception, the inference strategies that characterize visuo-vestibular integration in the presence of sensory conflict remain poorly understood (Dokka et al., 2015). For example, a recent study has found that observers predominantly integrated visual and vestibular cues even in the presence of large spatial discrepancies (de Winkel et al., 2015) – whereas a subsequent work has presented evidence in favor of causal inference (de Winkel et al., 2017). Furthermore, these studies did not vary cue reliability – a manipulation that is critical to test whether a Bayes-optimal inference strategy or a suboptimal approximation was used (Ma, 2012).

Another aspect that can influence the choice of inference strategy is the type of inference performed by the observer. In particular, de Winkel and colleagues (de Winkel et al., 2015, 2017) asked subjects to indicate the perceived direction of inertial heading – an ‘implicit’ inference task as subjects implicitly assessed the causal relationship between visual and vestibular cues on their way to indicate the final (integrated or segregated) heading percept. Even in the presence of spatial disparities as high as 90°, subjects fully integrated visual and vestibular cues (de Winkel et al., 2015; but see also de Winkel et al., 2017). It is plausible that performing an explicit inference task, which forces subjects to indicate whether visual and vestibular cues arose from the same or different events, may elicit different inference strategies, as previously reported in category-based induction (Chen, Ross, & Murphy, 2014), multi-cue judgment (Evans, 2008), and sensorimotor decision-making (Trommershäuser, Maloney, & Landy, 2008). While some studies have tested both explicit and implicit causal inference (Körding et al., 2007; Rohe & Noppeney, 2015b; Wallace et al., 2004), to our knowledge only one previous study contemplated the possibility of different strategies between implicit and explicit inference tasks (Rohe & Noppeney, 2015b), and a systematic comparison of inference strategies in the two tasks has never been carried out within a larger computational framework.

Thus, the goal of this work is two-fold. First, we introduce a set of techniques to perform robust, efficient Bayesian factorial model comparison of a variety of Bayesian and non-Bayesian models of causal inference in multisensory perception. Factorial comparison is a way to simultaneously test different orthogonal hypotheses about the observers (Acerbi, Vijayakumar, & Wolpert, 2014; Acerbi, Wolpert, & Vijayakumar, 2012; Rohe & Noppeney, 2015b; van den Berg, Awh, & Ma, 2014). Our approach is fully Bayesian in that we consider both parameter and model uncertainty, improving over previous analyses which used point estimates for the parameters and compared individual models. A full account of uncertainty in both parameter and model space, by marginalizing over parameters and model components, is particularly prudent when dealing with internal processes, such as decision strategies, which may have different latent explanations. An analysis that disregards such uncertainty might produce unwarranted conclusions about the internal processes that generated the observed behavior (Acerbi, Ma, & Vijayakumar, 2014). Second, we demonstrate our methods by quantitatively comparing the decision strategies underlying explicit and implicit causal inference in visuo-vestibular heading perception within the framework of Bayesian model comparison. We found that even though the study of explicit and implicit inference in isolation might suggest different inference rules, a joint analysis that combines all available evidence points to no difference between tasks, with subjects performing some form of causal inference in both the explicit and implicit tasks that used identical experimental setups.

In sum, we demonstrate how state-of-the-art techniques for model building, fitting, and comparison, combined with advanced analysis tools, allow us to ask nuanced questions about the observer’s decision strategies in causal inference. Importantly, these methods come with a number of diagnostics, sanity checks and a rigorous quantification of uncertainty that allow the experimenter to be explicit about the weight of evidence.

## Results

### Computational framework

We compiled a diverse set of computational techniques to perform robust Bayesian comparison of models of causal inference (CI) in multisensory perception, which we dub the ‘Bayesian cookbook for CI in multisensory perception’, or herein simply ‘the cookbook’. The main goal of the cookbook is to characterize observers’ decision strategies underlying causal inference, and possibly other details thereof, within a rigorous Bayesian framework that accounts for both parameter uncertainty and model uncertainty. The cookbook is ‘doubly-Bayesian’ in that it affords a fully Bayesian analysis of observers who may or may not be performing Bayesian inference themselves (Huszár, Noppeney, & Lengyel, 2010). Fully Bayesian model comparison is computationally intensive, hence the cookbook is concerned with efficient algorithmic solutions.

The cookbook comprises of: (a) techniques for fast evaluation of a large number of CI observer models; (b) procedures for model fitting via maximum likelihood, and approximating the Bayesian posterior of the parameters via Markov Chain Monte Carlo (MCMC); (c) state-of-the-art methods to compute model comparison metrics and perform factorial model selection. It is noteworthy that this cookbook is general and can be applied to multisensory perception across sensory domains.

### Causal inference in heading perception

We demonstrate our framework taking as a case study the comparison of explicit vs. implicit causal inference strategies in heading perception. In this section we briefly summarize our methods. Extended details and description of the cookbook can be found in the Methods and Appendix A, B, and C.

#### Experiments

Human observers were presented with synchronous visual (*s*_vis_) and vestibular (*s*_vest_) headings in the same direction (*C* = 1) or in different directions (*C* = 2) separated by a directional disparity Δ (Figure 1A). Mean stimulus direction (−25°,−20°,−15°,…,25°), cue disparity (0°, ±5°, ±10°, ±20°, and ±40°), and visual cue reliability *c_vis_* (coherence: high, medium and low) changed randomly on a trial-by-trial basis (Figure 1B). On each trial, non-zero disparity was either positive (vestibular heading to the right of visual heading) or negative. Observers (*n* = 11) first performed several sessions of an *explicit* causal inference task (‘unity judgment’), in which they indicated if the visual and vestibular stimuli signaled heading in the *same* direction (‘common cause’) or in *different* directions (‘different causes’). The same observers then participated in a number of sessions of the *implicit* causal inference task (‘inertial left/right discrimination’) wherein they indicated if their perceived inertial heading (vestibular) was to the left or right of straight forward. Both tasks consisted of a binary classification (same/different or left/right) with identical experimental apparatus and stimuli. No feedback was given to subjects about the correctness of their response. All observers also performed a number of practice trials and an initial session of a ‘unisensory left/right discrimination’ task in which they reported heading direction (left or right of straight forward) of visual or vestibular stimuli presented in isolation.

**Figure 1.**
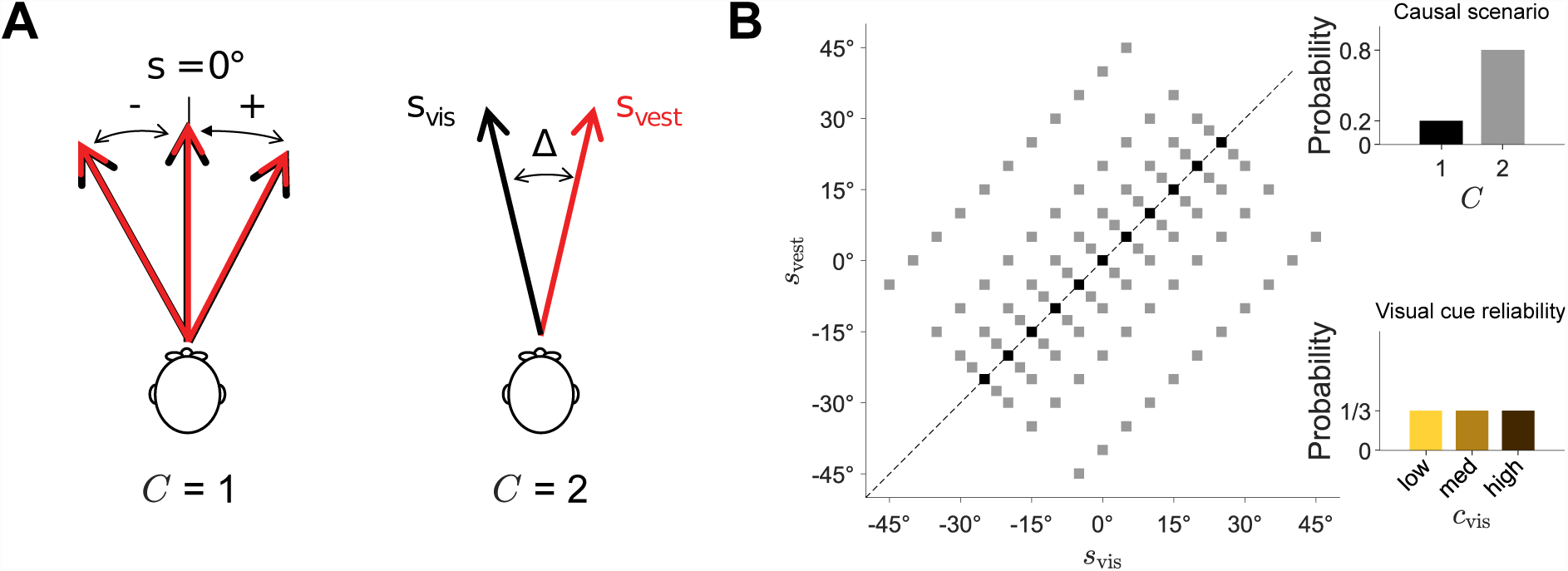
Experiment layout. **A:** Subjects were presented with visual (*s*_vis_) and vestibular (*s*_vis_) headings either in the same direction (*C* = 1) or in different directions (*C* = 2). In different sessions, subjects were asked to judge whether stimuli had the same cause (‘unity judgment’, explicit causal inference) or whether the vestibular heading was to the left or right of straight forward (‘inertial discrimination’, implicit causal inference). **B:** Distribution of stimuli used in the task. Mean stimulus direction was drawn from a discrete uniform distribution (−25°,−20°,−15°,…,25°). In 20% of the trials, *s*_vis_ ≡ *s*_vest_ (‘same’ trials, *C* = 1); in the other 80% (‘different’, *C* = 2), disparity was drawn from a discrete uniform distribution (±5°, ±10°, ±20°, ±40°), which led to a correlated pattern of heading directions *s*_vis_ and *s*_vest_. Visual cue reliability *c*_vis_ was also drawn randomly on each trial (high, medium and low).

#### Theory

For each task we built a set of observer models by factorially combining three model components that represent different assumptions about the observers: shape of sensory noise, type of prior over stimuli, and causal inference strategy (Figure 2A).

**Figure 2.**
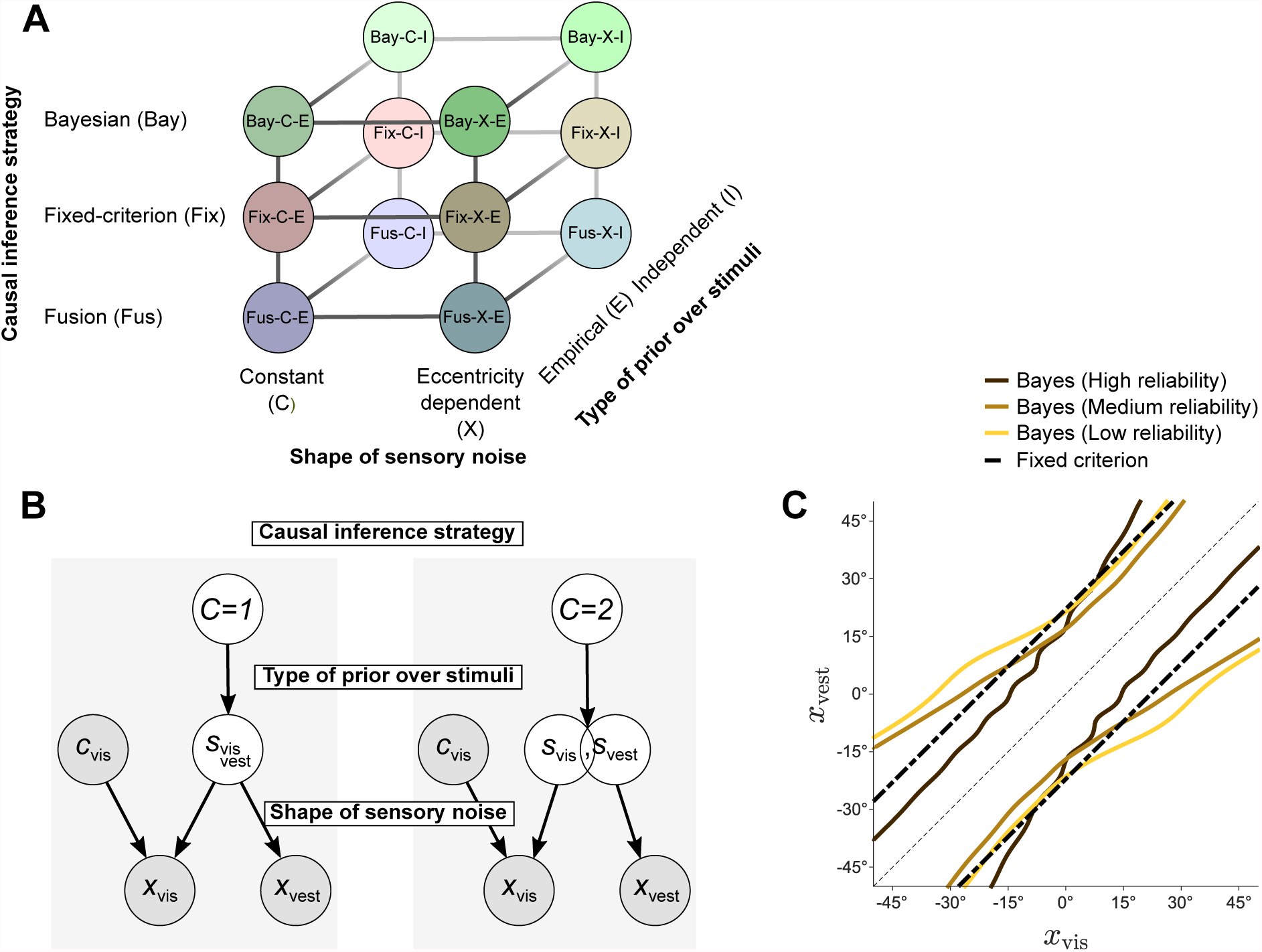
Observer models. **A:** Observer models were built by factorially combining three model components: Causal inference strategy, Shape of sensory noise, and Type of prior over stimuli (see text for details). Note that Fusion (‘Fus’) models are instantianted in different ways depending on the task. **B:** Graphical representation of the observer model. In the left panel (*C* = 1), the visual (*s*_vis_) and vestibular (*s*_vest_) heading direction have a single, common cause. In the right panel (*C* = 2), *s*_vis_ and *s*_vest_ have separate sources, although not necessarily statistically independent. The observer has access to noisy sensory measurements *x*_vis_, *x*_vest_, and knows the visual reliability level of the trial *c*_vis_. The observer is either asked to infer the causal structure of the task, *C* ≟ 1 (unity judgment, explicit inference), or whether the vestibular stimulus is rightward of straight ahead, 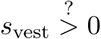 (inertial discrimination, implicit inference). Model factors affect different stages of the observer model: how the observer decides to combine the two causal scenarios (causal inference strategy); the prior over heading directions *p*_prior_(*s*_vis_*, s*_vest_|*C*) (type of prior over stimuli); and shape of noisy measurements distributions *p*(*x*_vis_|*s*_vis_*,c*_vis_) and *p*(*x*_vest_|*s*_vest_) (shape of sensory noise; note that this model factor affects equally both the generative model, that is how noisy measurements are generated, and the observer model, that is the observer’s beliefs about such noise). **C:** Example decision boundaries for the Bay-X-E model (for the three reliability levels), and for the Fixmodel, for a representative observer. The observer would report ‘unity’ when the noisy measurements *x*_vis_, *x*_vest_ fall within the boundaries. Note that the Bayesian decision boundaries expand with larger noise. Nonlinearities in the Bayesian boundaries are due to the interaction between eccentricity-dependence of the noise and the prior. In particular, wiggles are due to the discrete empirical prior.

In each trial of the explicit and implicit causal inference tasks, two stimuli are presented: a visual heading *s*_vis_ with known reliability *c*_vis_ ∊ {high, medium, low}, and a vestibular heading *s*_vest._ We assume that stimuli *s*_vis_, *s*_vest_ induce noisy measurements *x*_vis_ (resp., *x*_vest_) with conditionally independent distributions *p*(*x*_vis_|*s*_vis_ *c*_vis_) and *p*(*x*_vest_|*s*_vest_). For any stimulus *s* we assume that the noise distribution is Gaussian centered on *s* and with variance *σ*^2^(*s*). For each observer model we consider a variant in which *σ*^2^ depends only on the stimulus modality and reliability (*constant*, ‘C’) and a variant in which *σ*^2^ (*s*) also depends on stimulus location, growing with heading eccentricity, that is with the distance from 0° (*eccentricity-dependent*, ‘X’; see Methods). With a few notable exceptions (de Winkel et al., 2015, 2017; Odegaard et al., 2015), stimulus-dependence in the noise has been generally ignored in previous work (Beierholm et al., 2009; Körding et al., 2007; Rohe & Noppeney, 2015a, 2015b; Wozny et al., 2010). The base noise magnitude is governed by model parameters *σ*_0vest_ and *σ*_0vis_(*c*_vis_), where the latter is one parameter per visual reliability level. The eccentricity-dependent noise model has additional parameters *w*_vest_ and *w*_vis_ which govern the growth of noise with heading eccentricity (see Methods and Appendix A for details). We assume that the noise distribution equally affects both the generative model and the observer’s decision model, that is, observers have an approximately correct model of their own sensory noise (Alais & Burr, 2004; Ernst & Banks, 2002).

We assume that the observer considers two causal scenarios (Körding et al., 2007): either there is a single common heading direction (*C* = 1) or the two stimuli correspond to distinct headings (*C* = 2) (Figure 2B). If *C* = 1, the observer believes that the measurements are generated from the same underlying source *s* with prior distribution *p*_prior_(*s*). If *C* = 2, stimuli are believed to be distinct, but not necessarily statistically independent, with prior distribution *p*_prior_(*s*_vis_, *s*_vest_). For the type of these priors, we consider an *empirical* (‘E’) observer whose priors correspond to an approximation of the discrete, correlated distribution of stimuli in the task (as per Figure 1B); and an *independent* (‘I’) observer who uses a common and independent uni-dimensional prior for the two stimuli. Parameter *σ*_prior_ represents the SD of each independent prior (for ‘I’ priors), or of the prior over mean stimulus direction (for ‘E’ priors); whereas Δ_prior_ governs the SD of the prior over disparity (‘E’ priors only). See Methods for details.

We assume that observers are Bayesian in dealing with each causal scenario (*C* = 1 or *C* = 2), but may follow different strategies for weighting and combining information from the two causal hypotheses. Specifically, we consider three families of causal inference strategies. The Bayesian (‘Bay’) strategy computes the posterior probability of each causal scenario Pr(*C*|*x*_vis_, *x*_vest,_ *c*_vis_) based on all information available in the trial. The fixed-criterion (‘Fix’) strategy decides based on a fixed threshold of disparity between the noisy visual and vestibular measurements, disregarding reliability and other statistics of the stimuli. Finally, the fusion (‘Fus’) strategy disregards any location information, either always combining cues, or combining them with some probability (depending on whether the task involves implicit or explicit inference).

In the explicit inference task, the Bayesian (‘Bay’) observer reports a common cause if its posterior probability is greater than 0.5, Pr(*C* = 1|*x*_vis_, *x*_vest_, *c*_vis_) > 0.5. The prior probability of common cause, *p*_c_ ≡ Pr(*C* = 1), is a free parameter of the model. The fixed-criterion (‘Fix’) observer reports a common cause whenever the two noisy measurements are closer than a fixed distance *κ*_c_, that is |*x*_vis_ − *x*_vest_| < *κ*_c_, where the criterion *κ*_c_ is a free parameter that does not depend on stimulus reliability (Qamar et al., 2013). The fixed-criterion decision rule differs fundamentally from the Bayesian one in that it does not take cue reliability and other stimulus statistics into account (although noise will still affect behavior). As an example, Figure 2C shows the decision boundaries for the Bayesian (constant noise, empirical prior) and fixed-criterion rule for a representative observer. Finally, as a variant of the ‘fusion’ strategy we consider an observer that does not perform CI at all, but simply reports unity with probability *η*(*c*_vis_) regardless of stimulus disparity, where *η*_low_, *η*_med_, *η*_high_ are the only parameters of the model (*stochastic fusion*, ‘SFu’). This variant generalizes a trivial ‘forced fusion’ strategy (*η* ≡ 1) that would always report a common cause in the explicit inference.

For the implicit inference task, the observer first computes the posterior probability of rightward vestibular motion, Pr(*s*_vest_ > 0°|*x*_vest_, *x*_vis_, *c*_vis_, *C* = *k*) for the two causal scenarios, *k* = 1, 2. The Bayesian (‘Bay’) observer then reports ‘right’ if the posterior probability of rightward vestibular heading, averaged over the Bayesian posterior over causal structures, is greater than 0.5. The fixed-criterion (‘Fix’) observer reports ‘right’ if Pr(*s*_vest_ > 0°|*x*_vest_, *x*_vis_, *c*_vis_, *C* = *k*_fix_) > 0.5, where *k*_fix_ = 1 if |*x*_vis_ − *x*_vest_| < *κ*_c_, and *k*_fix_ = 2 otherwise. Finally, for the Fusion strategy we consider here the *forced fusion* (‘FFu’) observer, for which *C* ≡ 1. The forced fusion observer is equivalent to a Bayesian observer with *p*_c_ ≡ 1, and to a fixed-criterion observer for *κ*_c_ → ∞.

Observers also performed a unisensory left/right heading discrimination task, in which either a visual or vestibular heading was presented on each trial. In this case observers were modeled as standard Bayesian observers that respond ‘right’ if Pr(*s*_vis_ > 0°|*x*_vis_, *c*_vis_) > 0.5 for visual trials, and if Pr(*s*_vest_ > 0°|*x*_vest_) > 0.5 for vestibular trials. These data were used to constrain the joint model fits (see below).

For all observer models and tasks (except stochastic fusion in the explicit task), we considered a lapse probability 0 ≤ λ ≤ 1 of the observer giving a random response. Finally, we note that the Bayesian observer models considered in our main analysis perform Bayesian model averaging (the proper Bayesian strategy). At the end of the Results section we will also consider a ‘probability matching’ suboptimal Bayesian observer (Wozny et al., 2010).

#### Analysis strategy

Our analysis strategy consisted of first examining subjects’ behavior separately in the explicit and implicit tasks via model fitting and comparison. We then compared the model fits across tasks to ensure that model parameters were broadly compatible, allowing us to aggregate data from different tasks without changing the structure of the models. Finally, we re-analyzed observers’ performance by jointly fitting data from all three tasks (explicit causal inference, implicit causal inference, and unisensory heading discrimination), thereby combining all available evidence to characterize subjects’ decision making processes.

Given the large number of models and distinct datasets involved, we coded each model using efficient computational techniques at each step (see Methods for details).

We fitted our models to the data first via maximum likelihood estimation (MLE), and then via Bayesian estimation of the posterior over parameters using Markov Chain Monte Carlo (MCMC). Posteriors are an improvement over point estimates in that they allow us to incorporate uncertainty over individual subjects’ model parameters in our analysis, and afford computation of more accurate comparison metrics (see below).

We computed for each task, subject, and model the leave-one-out cross-validation score (LOO) directly estimated from the MCMC output (Vehtari, Gelman, & Gabry, 2015; reported in Appendix D). LOO has several advantages over other model selection metrics in that it takes parameter uncertainty into account and provides a more accurate measure of predictive performance (Vehtari, Gelman, & Gabry, 2016; see Discussion). We combined model evidence (LOO scores) from different subjects and models using a hierarchical Bayesian approach for group studies (Stephan, Penny, Daunizeau, Moran, & Friston, 2009). For each model component within the model factors of interest (noise, prior, and causal inference strategy), we reported as the main summary statistic of the analysis the protected exceedence probability 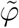, that is the (posterior) probability of a model component being the most likely component, above and beyond chance (Rigoux, Stephan, Friston, & Daunizeau, 2014). As a test of robustness, we also computed additional model comparison metrics: the corrected Akaike’s information criterion (AICc), the Bayesian information criterion (BIC), and an estimate of the log marginal likelihood (LML). While we prefer LOO as the main metric (see Discussion), we verified that the results of the model comparison were largely invariant of the choice of comparison metric.

Finally, for each model we estimated the absolute goodness of fit as the fraction of information gain above chance (where 0% is chance and 100% is the estimated intrinsic variability of the data, that is the entropy; Shen & Ma, 2016).

### Explicit inference task

We examined how subjects perceived the causal relationship of synchronous visual and vestibular headings as a function of disparity (*s*_vest_ − *s*_vis_, nine levels) and visual reliability level (high, medium, low; Figure 3A). Common cause reports were more frequent near zero disparities than for well-separated stimuli (Repeated-measures ANOVA with Greenhouse-Geisser correction; *F*_(1.82,18.17)_ = 76.0, *ϵ* = 0.23, *p* < 10^−4^, 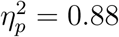). This means that observers neither performed complete integration (always reporting a common cause) nor complete segregation (never reporting a common cause). Common-cause reports were not affected by visual cue reliability alone (*F*_(1.23,12.33)_ = 1.84, *ϵ* = 0.62, *p* = .2, 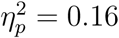), but were modulated by an interaction of visual reliability and disparity (*F*_(7.44,74.44)_ = 7.38, *ϵ* = 0.47, *p <* 10^−4^, 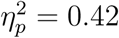). Thus, observers’ performance was affected by both cue disparity as well as visual cue reliability when explicitly reporting about the causal relationship between visual and vestibular cues. However, this does not necessarily mean that the subjects’ causal inference strategy took visual cue reliability into account. Changes in sensory noise may affect measured behavior even if the observer’s decision rule ignores such changes (Ma, 2012); a quantitative model comparison is needed to probe this question.

**Figure 3.**
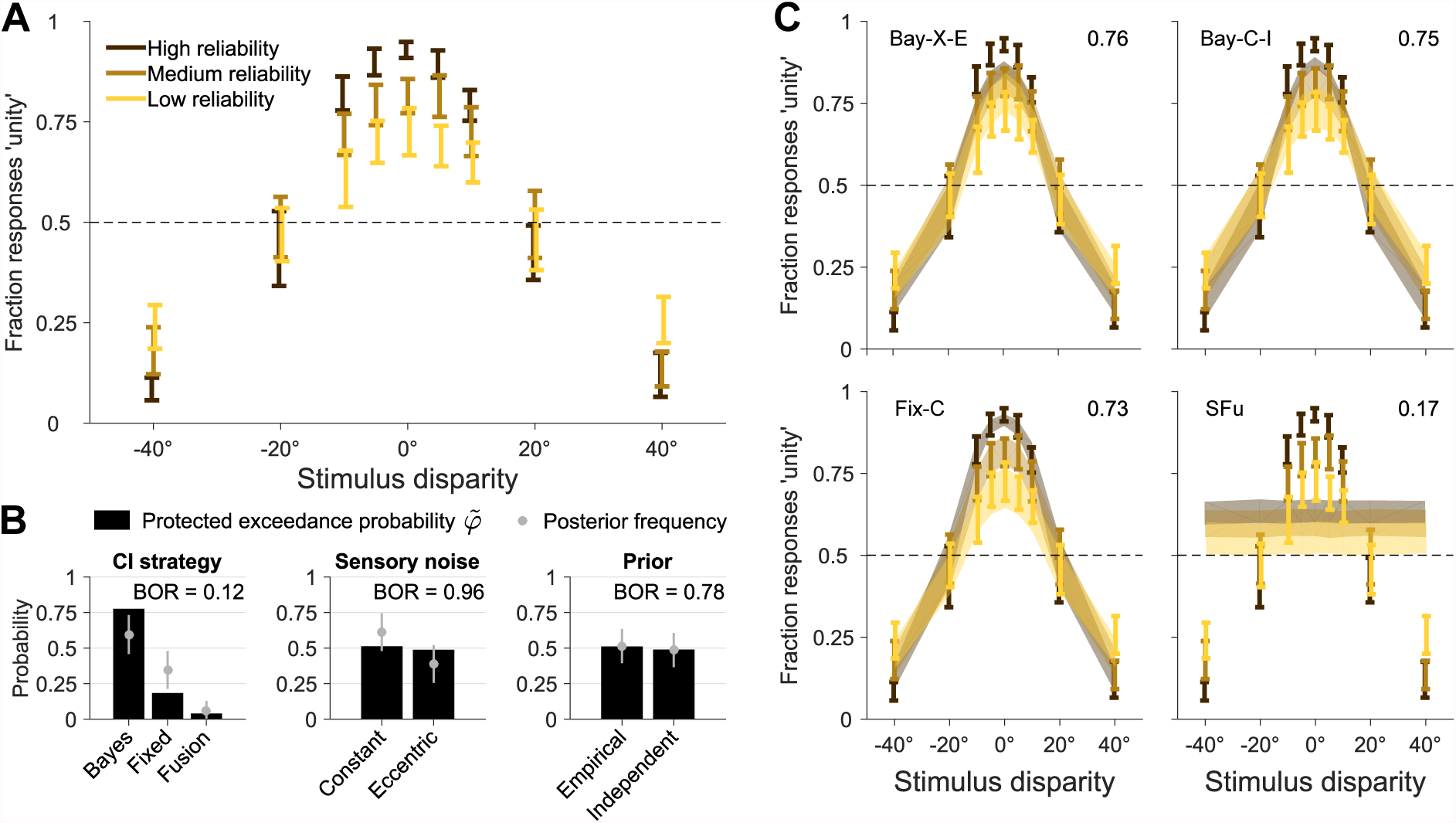
Explicit inference. Results of the explicit inference (unity judgment) task. **A:** Proportion of ‘unity’ responses, as a function of stimulus disparity (difference between vestibular and visual heading direction), and for different levels of visual cue reliability. Bars are ±1 SEM across subjects. Unity judgments are modulated by stimulus disparity and visual cue reliability. **B:** Protected exceedance probability 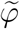 and estimated posterior frequency (mean ± SD) of distinct model components for each model factor. Each factor also displays the Bayesian omnibus risk (BOR). **C:** Model fits of several models of interest (see text for details). Shaded areas are ± SEM of model predictions across subjects. Numbers on top right of each panel report the absolute goodness of fit.

We compared a subset of models from the full factorial comparison (Figure 2A), since some models are equivalent when restricted to the explicit inference task. In particular, here fixed-criterion models are not influenced by the ‘prior’ factor, and the (stochastic) fusion model is not affected by sensory noise or prior, thus reducing the list of models to seven: Bay-C-E, Bay-C-I, Bay-X-E, Bay-X-I, Fix-C, Fix-X, SFu.

To assess the evidence for distinct determinants of subjects’ behavior, we combined LOO scores from individual subjects and models with a hierarchical Bayesian approach (Stephan et al., 2009; Figure 3B). Since we are investigating model factors that comprise of an unequal number of models, we reweighted the prior over models such that distinct components within each model factor had equal prior probability (Fixmodels had 2× weight, and SFu4×). In Figure 3B we report the protected exceedance probabilities 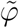 and, for reference, the posterior model frequencies they are based on, and the Bayesian omnibus risk (BOR), which is the estimated probability that the observed differences in factor frequencies may be due to chance (Rigoux et al., 2014). We found that the most likely factor of causal inference was the Bayesian model 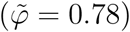, followed by fixed-criterion 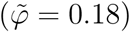 and probabilistic fusion 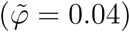. That is, fusion was ~ 24 times less likely to be the most representative model than any form of causal inference combined, which is strong evidence against fusion, and in agreement with our model-free analysis. The Bayesian strategy was ~ 3.5 times more likely than the others, which is positive but not strong evidence (Kass & Raftery, 1995). Conversely, the explicit inference data do not allow us to draw conclusions about noise models (constant vs. eccentric) or priors (empirical vs. independent), as we found that all factor components are about equally likely 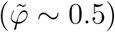.

At the level of specific models – as opposed to aggregate model factors –, we found that the probability of being the most likely model was almost equally divided between fixed-criterion (C-I) and Bayesian (either X-E or C-I). All these models yielded reasonable fits (Figure 3C), which captured a large fraction of the noise in the data (absolute goodness of fit ≈ 76% ± 3%; see Methods); a large improvement over a constant-probability model, which had a goodness of fit of 14 ± 5%. For comparison, we also show in Figure 3C the stochastic fusion model, which had a goodness of fit of 17% ± 5%. Visually, the Fixmodel in Figure 3C seems to fit better the group data, but we found that this is an artifact of projecting the data on the disparity axis. Disparity is the only relevant dimension for the Fixmodel; whereas Baymodels fits the data along all dimensions. The visual superiority of the Fixmodel wanes when the data are visualized in their entirety (see Appendix E).

We verified robustness of our findings by performing the same hierarchical analysis with different model comparison metrics. All metrics were in agreement with respect to the Bayesian causal inference strategy as the most likely, and the same three models being most probable (although possibly with different ranking). BIC and marginal likelihood differed from LOO and AICc mainly in that they reported a larger probability for the constant vs. eccentricity-dependent noise (probability ratio ~ 4.6, which is positive but not strong evidence).

These results combined provide strong evidence that subjects in the explicit inference task took into account some elements of the statistical structure of the trial (disparity, and possibly cue reliability) to report unity judgments, consistent with causal inference, potentially in a Bayesian manner. From these data, it is unclear whether observers took into account the empirical distribution of stimuli, and whether their behavior was affected by eccentricity-dependence in the sensory noise.

### Implicit inference task

We examined the bias in the reported direction of inertial heading computed as (minus) the point of subjective equality for left/rightward heading choices (L/R PSE), for each visual heading and visual cue reliability (Figure 4A). A shift in the vestibular L/R PSE away from zero represents a bias of the estimated inertial heading in the opposite direction. The bias was significantly affected by visual heading (Repeated-measures ANOVA; *F*_(0.71,7.08)_ = 19.67, *ϵ* = 0.07, *p* = .004, 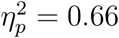). We found no main effect of visual cue reliability alone (*F*_(0.85,8.54)_ = 0.51, *ϵ* = 0.43, *p* = .47, 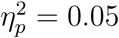), but there was a significant interaction of visual cue reliability and heading (*F*_(2.93,29.26)_ = 7.36, *ϵ* = 0.15, *p* < 10^−3^, 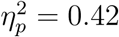). These data suggest that subjects’ perception of vestibular headings was modulated by visual cue reliability and visual stimulus, in agreement with previous work in visual-auditory localization (Rohe & Noppeney, 2015b). However, quantitative model comparison is required to understand the mechanism in detail since different processes could lead to similar patterns of observed behavior.

**Figure 4.**
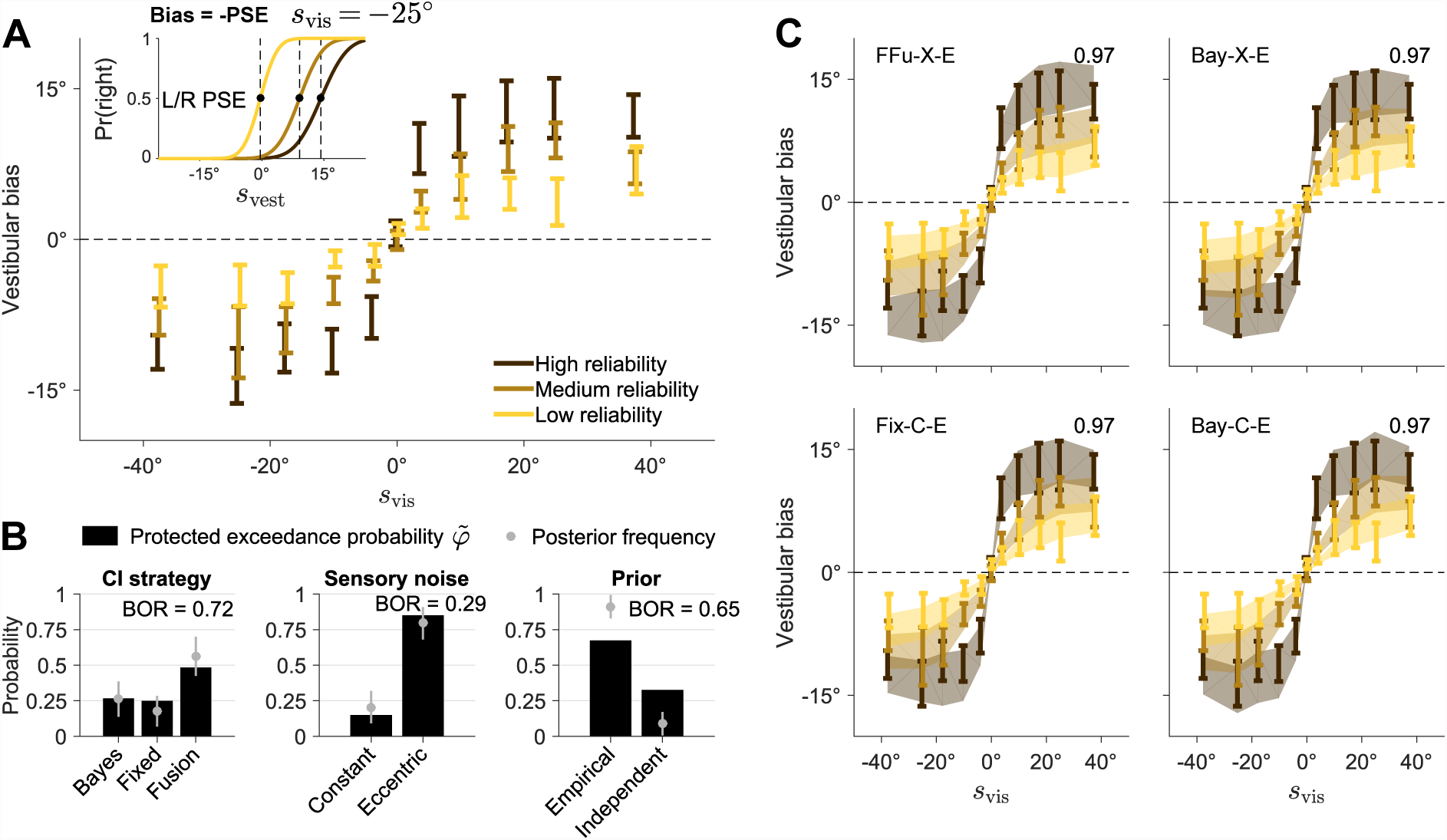
Implicit inference. Results of the implicit inference (left/right inertial discrimination) task. **A:** Vestibular bias as a function of co-presented visual heading direction *s*_vis_, at different levels of visual reliability. Bars are ±1 SEM across subjects. The inset shows a cartoon of how the vestibular bias is computed as minus the point of subjective equality of the psychometric curves of left/right responses (L/R PSE) for vestibular stimuli *s*_vest_, for a representative subject and for a fixed value of *s*_vis_. The vestibular bias is strongly modulated by *s*_vis_ and its reliability. **B:** Protected exceedance probability 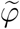 and estimated posterior frequency (mean ± SD) of distinct model components for each model factor. Each factor also displays the Bayesian omnibus risk (BOR). **C:** Model fits of several models of interests (see text for details). Shaded areas are ±1 SEM of model predictions across subjects. Numbers on top right of each panel report the absolute goodness of fit.

We performed a factorial comparison with all models in Figure 2A. In this case, factorial model comparison via LOO was unable to uniquely identify the causal inference strategy adopted by observers (Figure 4B). Forced fusion was slightly favored 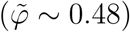, followed by Bayes 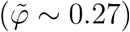 and fixed-criterion 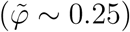, suggesting that all strategies were similar to forced fusion. Conversely, eccentricity-dependent noise was found to be more likely than constant noise (ratio ~ 5.7), which is positive but not strong evidence, and empirical priors were marginally more likely than independent priors (~ 2.1). The estimated Bayesian omnibus risk was high (BOR ≥ 0.29), hinting at a large degree of similarity within all model factors such that observed differences could have arisen by chance.

All metrics generally agreed on the lack of evidence in favor of any specific inference strategy (with AICc and BIC tending to marginally favor fixed-criterion instead of fusion), and on empirical priors being more likely. As a notable difference, marginal likelihood and BIC reversed the result about noise models, favoring constant noise models over eccentricity-dependent ones.

In terms of individual models, the most likely models according to LOO were, in order, forced fusion (X-E), Bayesian (X-E), and fixed-criterion (C-E). However, other metrics also favored other models; for example, Bayesian (C-E) was most likely according to the marginal likelihood. All these models obtained similarly good fits to individual data (Figure 4C; absolute goodness of fit ≈ 97%). For reference, a model that responds ‘rightward motion’ with constant probability performed about at chance (goodness of fit ≈ 0.3 ± 0.1%).

In sum, our analysis shows that the implicit inference data alone are largely inconclusive, possibly because almost all models behave similarly to forced fusion. To further explore our results, we examined the posterior distribution of the prior probability of common cause parameter *p*_c_ across Bayesian models, and of the criterion *κ*_c_ for fixed-criterion models (Figure 5, bottom left panels). In both cases we found a broad distribution of parameters, with only a mild accumulation towards ‘forced fusion’ values (*p*_c_ = 1 or *κ*_c_ ≳ 90°), suggesting that subjects were not completely performing forced fusion. Thus, it is possible that by constraining the inference with additional data we would be able to draw more defined conclusions.

**Figure 5.**
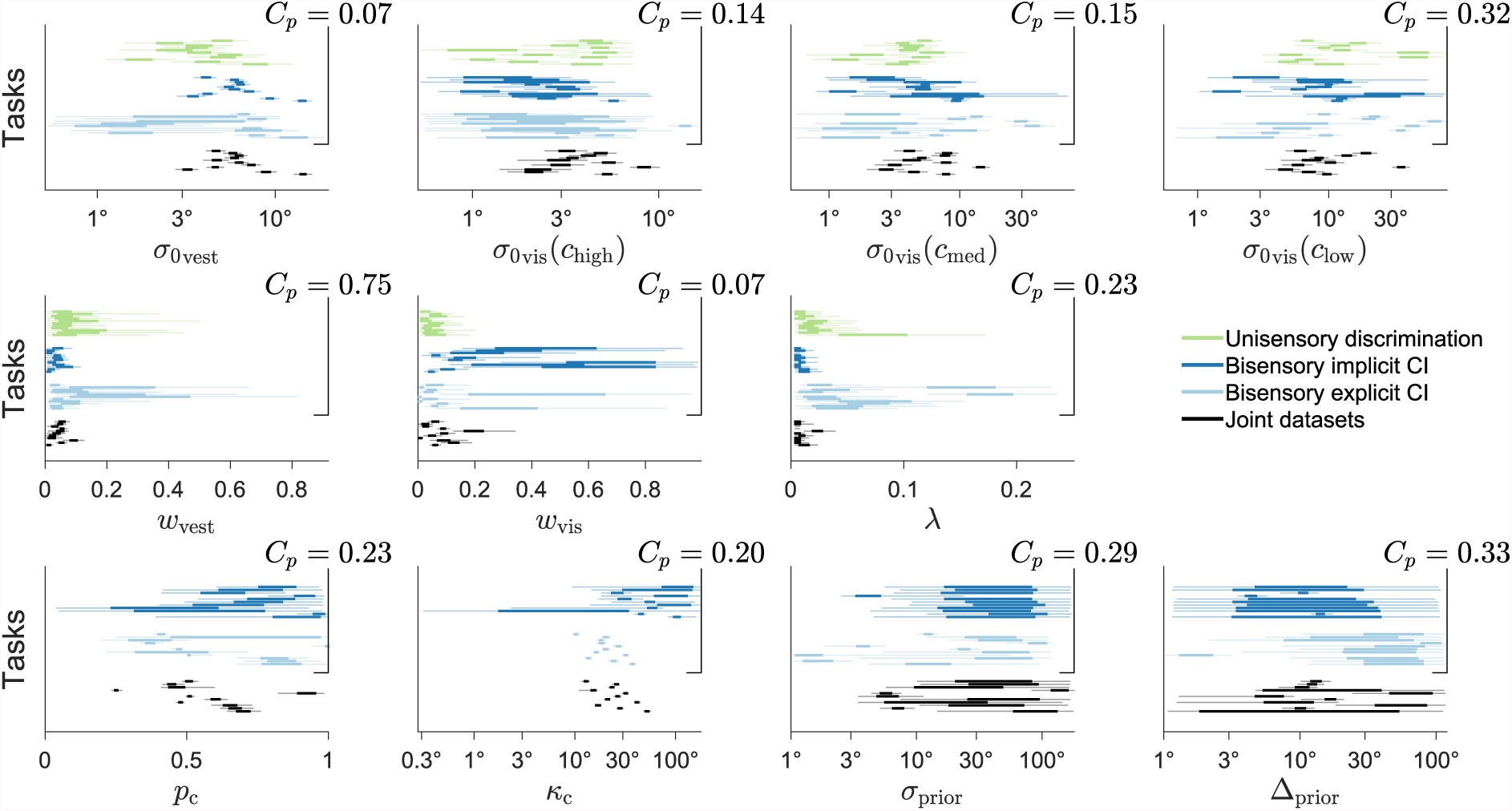
Posteriors over model parameters. Each panel shows the marginal posterior distributions over a single parameter for each subject and task. Each line is an individual subject’s posterior (thick line: interquartile range; light line: 95% credible interval); different colors correspond to different tasks. For each subject and task, posteriors are marginalized over models according to their posterior probability (see Methods). For each parameter we report the across-tasks compatibility probability *C_p_*, that is the (posterior) probability that subjects were best described by the assumption that parameter values were the same across separate tasks, above and beyond chance. The first two rows of parameters compute compatibility across all three tasks, whereas in the last row compatibility only includes the bisensory tasks (bisensory inertial discrimination and unity judgment), as these parameters are irrelevant for the unisensory task.

### Joint Model Fits

Data from the explicit and implicit causal inference tasks, when analyzed separately, afforded only weak conclusions about subjects’ behavior. The natural next step is to combine datasets from the two tasks along with the data from the unisensory heading discrimination task in order to better constrain the model fits.

Before performing such joint fit, we verified whether there was evidence that model parameters changed substantially across tasks, in which case we might have had to change the structure of the models (e.g., by introducing a subset of distinct parameters for different tasks; Acerbi, Vijayakumar, & Wolpert, 2014). For each model parameter, we computed the across-tasks compatibility probability *C_p_* (Figure 5), which is the (posterior) probability that subjects were most likely to have the same parameter values across tasks, as opposed to different parameters, above and beyond chance (see Methods for details). We found at most mild evidence towards difference of parameters across the three tasks, but no strong evidence (all *C_p_* > .05). Therefore, we proceeded in jointly fitting the data with the default assumption that parameters were shared across tasks.

For the joint fits there are nine possible models for the CI strategy (three explicit inference × three implicit inference strategies). However, we considered only a subset of plausible combinations, to avoid ‘model overfitting’ (see Discussion). First, we disregarded the stochastic fusion strategy for the explicit task, since this strategy was strongly rejected by the explicit task data alone. Second, if subjects performed some form of CI (Bayesian or fixed-criterion) in both tasks, we forced it to be the same. This reduces the model space for the causal inference strategy to four components: Bay/Bay, Fix/Fix, Bay/FFu, Fix/FFu(explicit/implicit task). Combined with the prior and sensory noise factors as per Figure 2A, this leads to sixteen models.

Factorial model comparison via LOO found that the most likely causal inference strategy was fixed-criterion 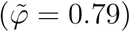, followed by Bayesian 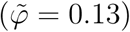, and then by forced fusion in the implicit task (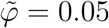 paired with Bayesian explicit inference, 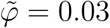 paired with fixed-criterion explicit inference; Figure 6A). This is positive evidence that subjects were performing some form of causal inference also in the implicit task, as opposed to mere forced fusion (ratio ~ 11.4). Moreover, we found strong evidence for eccentricity-dependent over constant noise 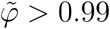, ratio ~ 132.7). Instead, the joint data were still inconclusive about the prior adopted by the subjects, with only marginal evidence for the empirical prior over the independent prior (~ 2.9).

In terms of specific models, the most likely model was fixed-criterion (X-E), followed by Bayesian (X-E), and explicit Bayesian / implicit forced fusion (both X-I and X-E). The best models gave a good description of the individual joint data, with an absolute goodness of fit of ≈ 91% ± 1% (Figure 6B).

**Figure 6.**
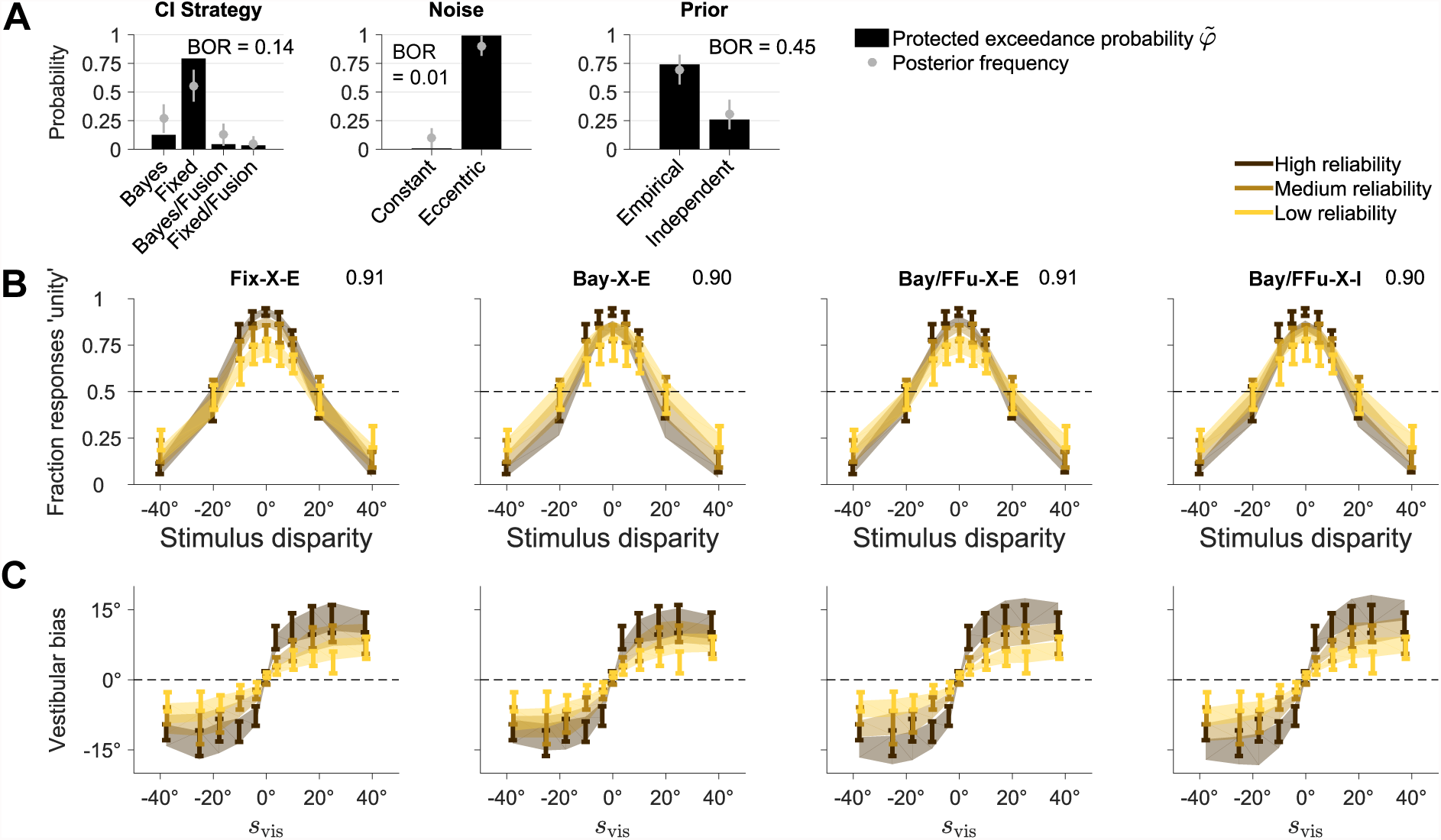
Joint fits. Results of the joint fits across tasks. **A:** Protected exceedance probability 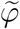 and estimated posterior frequency (mean ± SD) of distinct model components for each model factor. Each factor also displays the Bayesian omnibus risk (BOR). **B:** Joint model fits of the explicit inference (unity judgment) task, for different models of interest. Each panel shows the proportion of ‘unity’ responses, as a function of stimulus disparity and for different levels of visual reliability. Bars are ±1 SEM of data across subjects. Shaded areas are ±1 SEM of model predictions across subjects. Numbers on top right of each panel report the absolute goodness of fit across all tasks. **C:** Joint model fits of the implicit inference task, for the same models of panel B. Panels show vestibular bias as a function of co-presented visual heading direction *s*_vis_, and for different levels of visual reliability. Bars are ±1 SEM of data across subjects. Shaded areas are ±1 SEM of model predictions across subjects.

Examination of the subjects’ posteriors over parameters for the joint fits (Table 1 and Figure 5, black lines) showed reasonable results. The base visual noise parameters were generally monotonically increasing with decreasing visual cue reliability; the vestibular base noise was roughly of the same magnitude as the medium visual cue noise (as per experiment design); both visual and vestibular noise increased mildly with the distance from straight ahead; subjects had a small lapse probability. For Bayesian models, *p*_c_ was substantially larger than the true value, 0.20 (*t*-test *t*_(10)_ = 10.8, *p* < 10^−4^, *d* = 3.25), suggesting that observers generally thought that heading directions had a higher a priori chance to be the same. Nonetheless, for all but one subject *p*_c_ was far from 1, suggesting that subjects were not performing forced fusion either. An analogous result holds for the fixed criterion *κ*_c_, which was smaller than the largest disparity between heading directions. We found that prior parameters *σ*_prior_ and Δ_prior_ had a lesser impact on the models, and their exact values were less crucial, with generally wide posteriors.

**Table 1.** Joint fit parameters.

| Parameter | Description | Posterior mean | Allowed range |
| --- | --- | --- | --- |
| All tasks |  |  |  |
| $\sigma_{0\text{vest}}$ | Vestibular base noise | $6.49^\circ \pm 0.90^\circ$ | $[0.5^\circ, 80^\circ]^\dagger$ |
| $\sigma_{0\text{vis}}(c_{\text{high}})$ | Visual base noise (high coherence) | $4.08^\circ \pm 0.54^\circ$ | $[0.5^\circ, 80^\circ]^\dagger$ |
| $\sigma_{0\text{vis}}(c_{\text{med}})$ | Visual base noise (medium coherence) | $6.32^\circ \pm 1.00^\circ$ | $[0.5^\circ, 80^\circ]^\dagger$ |
| $\sigma_{0\text{vis}}(c_{\text{low}})$ | Visual base noise (low coherence) | $11.57^\circ \pm 2.67^\circ$ | $[0.5^\circ, 80^\circ]^\dagger$ |
| $w_{\text{vest}}$ | Vestibular noise eccentricity | $0.04 \pm 0.01$ | $[0, 1]$ |
| $w_{\text{vis}}$ | Visual noise eccentricity | $0.07 \pm 0.02$ | $[0, 1]$ |
| $\lambda$ | Lapse rate | $0.01 \pm 0.01$ | $[0, 1]$ |
| Bisensory only |  |  |  |
| $p_c$ | Prior of common cause (Baymodels) | $0.56 \pm 0.05$ | $[0, 1]$ |
| $\kappa_c$ | Fixed criterion (Fixmodels) | $26.50^\circ \pm 3.52^\circ$ | $[0.25^\circ, 180^\circ]^\dagger$ |
| $\sigma_{\text{prior}}$ | Central prior width | $49.77^\circ \pm 12.08^\circ$ | $[1^\circ, 120^\circ]^\dagger$ |
| $\Delta_{\text{prior}}$ | Disparity prior width | $23.51^\circ \pm 6.39^\circ$ | $[1^\circ, 120^\circ]^\dagger$ |

Posterior means of parameters in the joint fit, marginalized over models according to each subject’s posterior model probability, and averaged across subjects ( ± SEM). For reference, we also report the parameter range used for the optimization and MCMC sampling. † These parameters were transformed and fitted in log space.

Finally, we verified that our results did not depend on the chosen comparison metric. Remarkably, the findings regarding causal inference factors were quantitatively the same for all metrics, demonstrating robustness of our main result. Marginal likelihood and BIC differed from LOO and AICc in that they only marginally favored eccentricity-dependent noise models, showing that conclusions over the noise model may depend on the specific choice of metric. All metrics agreed in marginally preferring the empirical prior over the independent prior.

In conclusion, when combining evidence from all available data, our model comparison shows that subjects were most likely performing some form of causal inference instead of forced fusion, for both the explicit and the implicit CI tasks. In particular, we find that a fixed-criterion, non-probabilistic decision rule (i.e., one that does not take uncertainty into account) describes the joint data better than the Bayesian strategy, although with some caveats (see Discussion).

### Sensitivity analysis and model validation

Performing a factorial comparison, like any other statistical analysis, requires a number of somewhat arbitrary choices, loosely motivated by previous studies, theoretical considerations, or a preliminary investigation of the data (being aware of the ‘garden of forking paths’; Gelman & Loken, 2013). As good practice, we want to check that our main findings are robust to changes in the setup of the analysis, or be able to report discrepancies.

We take as our main result the protected exceedance probabilties 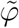 of the model factors in the joint analysis (Figure 6A, reproduced in Figure 7, top row). In the following, we examine whether this finding holds up to several manipulations of the analysis framework.

**Figure 7.**
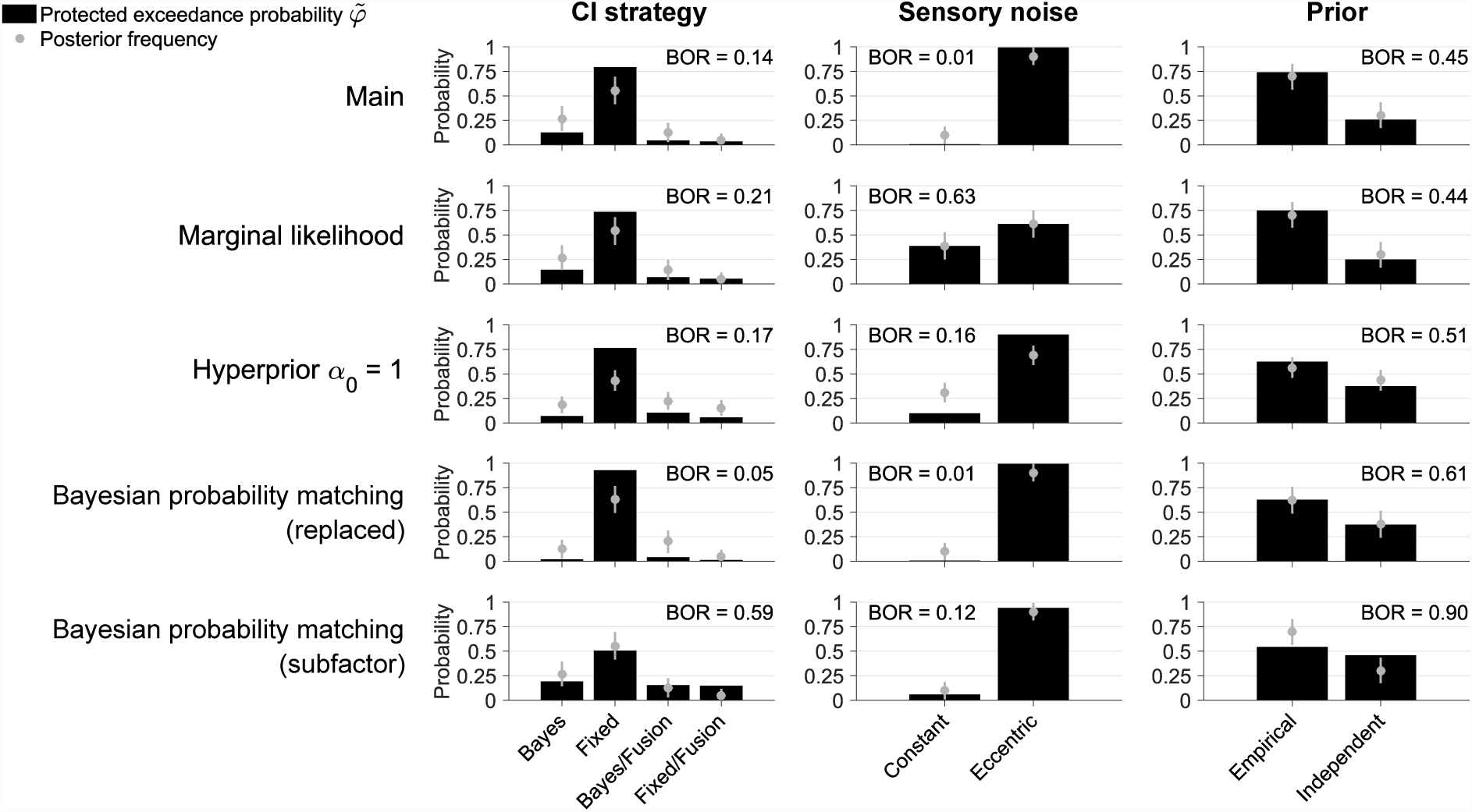
Sensitivity analysis of factorial model comparison. Protected exceedance probability 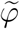 of distinct model components for each model factor in the joint fits. Each panel also shows the estimated posterior frequency (mean ± SD) of distinct model components, and the Bayesian omnibus risk (BOR). Each row represents a variant of the factorial comparison. 1st row: Main analysis (as per Figure 6A). 2nd row: Uses marginal likelihood as model comparison metric. 3rd row: Uses hyperprior *α*_0_ = 1 for the frequencies over models in the population (instead of a flat prior over model factors). 4th row: Uses ‘probability matching’ strategy for the Bayesian CI model (replacing model averaging). 5th row: Includes probability matching as a sub-factor of the Bayesian CI family (in addition to model averaging).

A first check consists of testing different model comparison metrics. In the previous sections, we have reported results for different metrics, finding in general only minor differences from our results obtained with LOO. As an example, we show here the model comparison using as metric an estimate of the marginal likelihood – the probability of the data under the model (Figure 7, 2nd row). We see that the marginal likelihood results agree with our results with LOO except for the sensory noise factor (see Discussion). Therefore, our conclusions about the CI strategy are not affected.

Second, the hierarchical Bayesian Model Selection method requires to specify a prior over frequencies of models in the population (Stephan et al., 2009). This (hyper)prior is specified via the concentration parameter vector ***α***_0_ of a Dirichlet distribution over model frequencies. For our analysis, since we focused on the factorial aspect, we chose an approximately ‘flat’ prior across model factors (see Methods for details), instead of the default flat prior over individual models (*α*_0_ = 1). We found that performing the group analysis with *α*_0_ = 1 did not change our results (Figure 7, 3rd row).

Another potential source of variation is specific model choices, or inclusion of model factors. For example, a common successful variant of the Bayesian CI strategy is ‘probability matching’, according to which the observer chooses the causal scenario (*C* = 1 or *C* = 2) randomly, proportionally to its posterior probability (Wozny et al., 2010). As a first check, we performed the model comparison again using a ‘probability matching’ Bayesian observer *instead* of our main ‘model averaging’ observer (Figure 7, 4th row). Results are similar to the main analysis. If anything, the fixed-criterion CI strategy gains additional evidence here, suggesting that probability matching is a worse description of the data than our original Bayesian CI model (as confirmed by looking at differences in LOO scores of individual subjects, e.g. for the Bay-X-E model; mean ± SEM: ΔLOO = −17.3 ± 5.7). A recent study in audio-visual causal inference perception has similarly found that probability matching provided a poor explanation of the data (Rohe & Noppeney, 2015b).

In the factorial framework we could also have performed the previous analysis in a different way, by considering ‘probability matching’ as a sub-factor of the Bayesian strategy, *together* with ‘model averaging’. As we have done before for the explicit inference task, we reassign prior probabilities to the models so that they are constant for each factor (in this case, the two Bayesian strategies get a 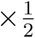 multiplier). Results of this alternative approach show an increase of evidence for the Bayesian CI family (7, bottom row). The values of 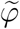 for the fusion models are also slightly higher, which is due to an increase of the Bayesian omnibus risk (the probability that the observed differences in factor frequencies are due to chance, a warning sign that there are too many models for the available data). This result and other lines of reasoning suggest caution when model factors contain an uneven number of models (see Discussion). Nonetheless, the main conclusion does not qualitatively change, in that observers performed some form of CI as opposed to forced fusion.

Finally, we performed several sanity checks, including a model recovery analysis to ensure the integrity of our analysis pipeline and that models of interest were meaningfully distinguishable (see Methods and Appendix B for details).

In conclusion, we have shown how the computational framework of Bayesian factorial model comparison, which is made possible by a combination of methods described in the cookbook, allows to explore multiple questions about aspects of subjects’ behavior in multisensory perception, and to account for uncertainty at different levels of the analysis in a principled, robust manner.

## Discussion

We presented a ‘cookbook’ of algorithmic recipes for robust Bayesian evaluation of observer models of causal inference that have widespread applications to multisensory perception and modeling perceptual behavior in general. We applied these techniques to investigate the decision strategies that characterize explicit and implicit causal inference in multisensory heading perception. Examination of observers’ behavior in the explicit and implicit inference tasks provided evidence that observers did not simply fuse visual and vestibular cues. Instead, observers integrated the multi-sensory cues based on their relative disparity, a signature of causal inference. Importantly, our framework affords investigation of whether humans adopt a statistically optimal Bayesian strategy or instead implement a heuristic decision rule which does not fully consider the uncertainty associated with the stimuli.

### Causal inference in multisensory heading perception

Our findings in the explicit inference task demonstrate that subjects used information about the discrepancy between the visual and vestibular cues to infer the causal relationship between them. Results in the implicit inference task alone were mixed, in that we could not clearly distinguish between alternative strategies, including forced fusion – in agreement with a previous finding (de Winkel et al., 2015). However, when we combined evidence from all tasks, we found that some form of causal inference was more likely than mere forced fusion, in agreement with a more recent study (de Winkel et al., 2017). Our findings suggest that multiple sources of evidence (e.g., different tasks) can help disambiguate causal inference strategies which might otherwise produce similar patterns of behavioral responses.

Our Bayesian analysis allowed us to examine the distribution of model parameters, in particular the causal inference parameters *p*_c_ and *κ*_c_, which govern the tendency to bind or separate cues for, respectively, a Bayesian and a heuristic fixed-criterion strategy. Evidence from all tasks strongly constrained these parameters for each subject. Interestingly, for the Bayesian models we found an average *p*_c_ much higher than the true experimental value (inferred *p*_c_ ~ 0.5 vs. experimental *p*_c_ = 0.2). This suggests that subjects had a tendency to integrate sensory cues substantially more than what the statistics of the task would require. Note that, instead, a Bayesian observer would be able to learn the correct value of *p*_c_ from noisy observations, provided some knowledge of the structure of the task. Our finding is in agreement with previous studies which demonstrated an increased tendency to combine discrepant visual and vestibular cues (Butler et al., 2010; Campos, Siegle, Mohler, Bülthoff, & Loomis, 2009; de Winkel et al., 2015; Kaliuzhna, Prsa, Gale, Lee, & Blanke, 2015; Prsa et al., 2012) and also a large inter-subject variability in *p*_c_, and not obviously related to the statistics of the task (Odegaard & Shams, 2016; Odegaard, Wozny, & Shams, 2017). We note that, in all studies so far, the ‘binding tendency’ (*p*_c_ or *κ*_c_) is a descriptive parameter of causal inference models that lacks an independent empirical correlate (as opposed to, for example, noise parameters, which can be independently measured). Understanding the origin of the binding tendency, and which experimental manipulations is sensitive to, is venue for future work (Odegaard & Shams, 2016; Odegaard et al., 2017).

Previous work has performed a factorial comparison of only causal inference strategies (Rohe & Noppeney, 2015b). Our analysis extends that work by including as latent factors the shape of sensory noise (and, thus, likelihoods) and type of priors (Acerbi, Vijayakumar, & Wolpert, 2014; Acerbi et al., 2012). Models in our set include a full computation of the observers’ posterior beliefs based on eccentricity-dependent likelihoods, which was only approximated in previous studies that considered eccentricity-dependence (de Winkel et al., 2015, 2017; Odegaard et al., 2015). Indeed, in agreement with a recent finding, we found an important role of eccentricity-dependent noise (Odegaard et al., 2015). Conversely, our analysis of priors was inconclusive, as our datasets were unable to tell whether people learnt the empirical (correlated) prior, or made an assumption of independence.

Our main finding, relative to the causal inference strategy, is that subjects performed causal inference both in the explicit and implicit tasks. Interestingly, from our analyses the most likely causal inference strategy is a fixed-criterion strategy, which crucially differs from the Bayesian strategy in that it does not take cue reliability into account – let alone optimally. This finding is seemingly at odds with a long list of results in multisensory perception, in which people are shown to take cue uncertainty into account (Butler et al., 2010; Ernst & Bülthoff, 2004; Fetsch et al., 2009; Gu et al., 2008). We note that this is not necessarily in contrast with existing literature, for several reasons. First, this result pertains specifically to the causal inference part of the observer model, and not how cues are combined once a common cause has been inferred (Rohe & Noppeney, 2015b). To our knowledge, no study has tested Bayesian models of causal inference against heuristic models that take into account disparity but not reliability, as it has been done for example in visual search (Ma, Navalpakkam, Beck, Van Den Berg, & Pouget, 2011; Shen & Ma, 2016) and visual categorization (Adler & Ma, 2016; Qamar et al., 2013). A quantitative modeling approach is needed – qualitatively analyzing the differences in behavior at different levels of reliability is not sufficient to establish that observers take uncertainty into account; patterns of observed differences may be due to a change in sensory noise even if the observer’s decision rule disregards cue reliability. Second, our results are not definitive – the evidence for fixed-criterion vs. Bayesian is positive but not decisive. Our interpretation of this result is that subjects are following some suboptimal decision rule which happens to be closer to fixed-criterion than to the Bayesian strategy for the presented stimuli and range of tested reliability levels. It is possible that with a wider range of stimuli and reliabilities, and possibly with different ways of reporting (e.g., estimation instead of discrimination), we would be able to distinguish the Bayesian strategy from a fixed-criterion heuristic.

Finally, we note that model predictions of our Bayesian models are good but still show systematic discrepancies from the data for the explicit inference task (Figs 3C and 6B). Previous work has found similar discrepancies in model fits of unity judgments data across multiple sensory reliabilities (e.g., see Figure 2A in Rohe & Noppeney, 2015b). This suggests that there is some element of model mismatch in current Bayesian causal inference models, possibly due to difference in noise models or to other processes that affect causal inference across cue reliabilities, which deserves further investigation.

### Bayesian factorial comparison

We performed our analysis within a factorial model comparison framework (van den Berg et al., 2014). Even though we were mainly interested in a single factor (causal inference strategy), previous work has shown that the inferred observer’s decision strategy might depend on other aspects of the observer model, such as sensory noise or prior, due to nontrivial interactions of all these model components (Acerbi, Ma, & Vijayakumar, 2014). Our method, therefore, consisted of performing inference across a family of observer models that explicitly instantiated plausible model variants. We then marginalized over details of specific observer models, looking at posterior probabilities of model factors, according to a hierarchical Bayesian model selection (BMS) approach (Rigoux et al., 2014; Stephan et al., 2009). We applied a few tweaks to the BMS method to account for our focus on factors as opposed to individual models (see Methods).

Our approach was fully Bayesian in that we took into account parameter uncertainty (by computing a metric, LOO, based on the full posterior distribution) and model uncertainty (by marginalizing over model components). A fully Bayesian approach has the advantages of explicitly representing uncertainty in the results (e.g., credible intervals over parameters), and of reducing the risk of overfitting, although it is not immune to it (Piironen & Vehtari, 2016).

In our case, we marginalized over models to reduce the risk of model overfitting, which is a complementary problem to parameter overfitting. Model overfitting is likely to happen when model selection is performed within a large number of discrete models. In fact, some authors recommend to skip discrete model selection altogether, preferring instead inference and Bayesian parameter estimation in a single overarching or ‘complete’ model (Gelman et al., 2013). We additionally tried to reduce the risk of model overfitting by balancing prior probabilities across factors, although we noted that this may not be enough to counterbalance the additional flexibility that a model factor gains by having more sub-models than a competitor. Our practical recommendation, until more sophisticated comparison methods are available, is to ensure that all model components within a factor have the same number of models, and to limit the overall number of models.

Our approach was also factorial in the treatment of different tasks, in that first we analyzed each bisensory task in isolation, and then combined trials from all data in a joint fit. The fully Bayesian approach allowed us to compute posterior distributions for the parameters, marginalized over models (see Figure 5), which in turn made it possible to test whether model parameters were compatibile across tasks, via the ‘compatibility probability’ metric. The compatibility probability is an approximation of a full model comparison to test whether a given parameter is the same or should differ across different datasets (in this case, tasks), where we consider ‘sameness’ to be the default (simplyfing) hypothesis. We note that if the identity or not of a parameter across datasets is a main question of the study, its resolution should be addressed via a proper model comparison.

With the joint fits, we found that almost all parameters were well constrained by the data (except possibly for the parameters governing the observers’ priors, *σ*_prior_ and Δ_prior_). An alternative option to better constrain the inference for scarce data or poorly identified parameters is to use informative priors (as opposed to non-informative priors), or a hierarchical approach that assumes a common (hyper)prior to model parameters across subjects (Friston et al., 2016).

### Model comparison metrics

The general goal of a model comparison metric is to score a model for goodness of fit and somehow penalize for model flexibility. In our analysis we have used Pareto-smoothed importance sampling leave-one-out cross-validation (PSIS-LOO; Vehtari et al., 2016) as the main metric to compare models (simply called LOO in the other sections for simplicity). In fact, there is a large number of commonly used metrics, such as (corrected) Akaike’s information criterion (AIC(c); see Burnham & Anderson, 2003), Bayesian information criterion (BIC; see Burnham & Anderson, 2003), deviance information criterion (DIC) (Spiegelhalter, Best, Carlin, & Van Der Linde, 2002), widely applicable information criterion (WAIC; Watanabe, 2010, and marginal likelihood (MacKay, 2003). The literature on model comparison is vast and with different schools of thought – by necessity here we only summarize some remarks. The first broad distinction between these metrics is between predictive metrics (AIC(c), DIC, WAIC, and PSIS-LOO; see Gelman, Hwang, & Vehtari, 2014), that try to approximate out-of-sample predictive error (that is, model performance on unseen data), and BIC and marginal likelihood, which try to establish the true model generating the data (MacKay, 2003). Another orthogonal distinction is between metrics based on point estimates (AIC(c) and BIC) vs. metrics that use partial to full information about the model’s uncertainty landscape (DIC, WAIC, PSIS-LOO, based on the posterior, and the marginal likelihood, based on the likelihood integrated over the prior).

First, when computationally feasible we prefer uncertainty-based metrics to point estimates, since the latter are only crude asymptotic approximations that do not take the model and the data into account, besides simple summary statistics (number of free parameters and possibly number of data points). Due to their lack of knowledge of the actual structure of the model, AIC(c) and BIC can grossly misestimate model complexity (Gelman et al., 2014).

Second, we have a ordered preference among predictive metrics, that is PSIS-LOO ≻ WAIC ≻ DIC ≻ AIC(c) (Gelman et al., 2014). The reason is that all of these metrics more or less asymptotically approximate full leave-one-out cross validation, with increasing degree of accuracy from right to left (Gelman et al., 2014; Vehtari et al., 2016). As mentioned before, AIC(c) works only in the regime of a large amount of data. DIC, albeit commonly used, has several issues and requires the posterior to be multivariate normal, or at least symmetric and unimodal – gross failures can happen when this is not the case, since DIC bases its estimate of model complexity on the mean (or some other measure of central tendency) of the posterior (Gelman et al., 2014). WAIC is a great improvement over DIC and does not require normality of the posterior, but its approximation is generally superseded by PSIS-LOO (Vehtari et al., 2016). Moreover, PSIS-LOO has a natural diagnostic, the exponents of the tails of the fitted Pareto distribution, which allows the user to know when the method may be in trouble (Vehtari et al., 2016). Full leave-one-out cross validation is extremely expensive, but PSIS-LOO only requires the user to compute the posterior via MCMC sampling, with no additional cost with respect to DIC or WAIC. Similarly to WAIC, PSIS-LOO requires the user to store for each posterior sample the log likelihood *per trial*, which with modern computers represent a negligible storage cost.

The marginal likelihood, or Bayes factor (of which BIC is a poor approximation), is an alternative approach to quantify model evidence, related to computing the posterior probability of the models (MacKay, 2003). While this is a principled approach, it entails several practical and theoretical issues. First, the marginal likelihood is generally hard to compute, since it usually involves a complicated, high-dimensional integral of the likelihood over the prior (although this computation can be simplified for nested models; see Verdinelli & Wasserman, 1995). Here, we have applied a novel approximation method for the marginal likelihood following ideas delineated in Caldwell and Liu (2014); Robert, Wraith, Goggans, and Chan (2009), obtaining generally sensible values. However, more work is needed to establish the precision and applicability of such technique. Besides practical computational issues, the marginal likelihood, unlike other metrics, is sensitive to the choice of prior over parameters, in particular its range (Gelman et al., 2013). Crucially, and against common intuition, this sensitivity does not reduce with increasing amounts of data. A badly chosen (e.g., excessively wide) prior for a non-shared parameter might change the marginal likelihood of a model by several points, thus affecting model ranking. The open issue of prior sensitivity has led some authors to largely discard model selection based on the marginal likelihood (Gelman et al., 2013).

For these reasons, we chose (PSIS-)LOO as the main model comparison metric. As a test of robustness, we also computed other metrics and verified that our results were largely independent of the chosen metric, or investigated the reasons when it was not the case.

As a specific example, in our analysis we found that LOO and marginal likelihood (or BIC) generally agreed on all comparisons, except for the sensory noise factor. Unlike LOO, the marginal likelihood tended to prefer constant noise models as opposed to eccentricity-dependent models. Our explanation of this discrepancy is that for our tasks eccentricity-dependence provides a consistent but small improvement to the goodness of fit of the models, which can be overrided by a large penalty due to model complexity (BIC), or to the chosen prior over the eccentricity-dependent parameters (*w*_vis_, *w*_vest_), whose range was possibly wider than needed (see Figure 5). The issue of prior sensitivity (specifically, dependence of results on an arbitrarily chosen range) can be attenuated by adopting a Bayesian hierarchical approach over parameters (or a more computationally feasibile approximation, known as empirical Bayes), which is venue for future work.

### Computational framework

Model evaluation, especially from a Bayesian perspective, is a time-consuming business. For this reason, we have compiled several state-of-the-art methods for model building, fitting and comparison, and made our code available.

The main issue of many common observer models in perception is that the expression for the (log) likelihood is not analytical, requiring numerical integration or simulation. To date, this limits the applicability of modern model specification and analysis tools, such as probabilistic programming languages, that exploit auto-differentiation and gradient-based sampling methods (e.g., Stan; Carpenter et al., 2016; or PyMC3; Salvatier, Wiecki, & Fonnesbeck, 2016). The goal of such computational frameworks is to remove the burden and technical details of evaluating the models from the shoulders of the modeler, who only needs to provide a model specification.

In our case, we strive towards a more modest goal of providing black-box algorithms for optimization and MCMC sampling that exhibit a larger degree of robustness than standard methods. In particular, for optimization (maximum likelihood estimation) we recommend Bayesian adaptive direct search (BADS; Acerbi & Ma, 2017), a technique based on Bayesian optimization (Jones, Schonlau, & Welch, 1998; Shahriari, Swersky, Wang, Adams, & de Freitas, 2016), which exhibits robustness to noise and jagged likelihood landscapes, unlike common optimization methods such as fminsearch (Nelder-Mead) and fmincon in MATLAB. Similarly, for MCMC sampling we propose a sampling method that combines the robustness and self-adaptation of slice sampling (Neal, 2003) and ensemble-based methods (Gilks, Roberts, & George, 1994). Crucially, our proposed method almost completely removes the need of expensive trial-and-error tuning on the part of the modeler, possibly one of the main reasons why MCMC methods and full evaluation of the posterior are relatively uncommon in the field (to our knowledge, this is the first study of causal inference in multisensory perception to adopt a fully Bayesian approach).

Our framework is similar to the concept behind the VBA toolbox, a MATLAB toolbox for probabilistic treatment of nonlinear models for neurobiological and behavioral data (Daunizeau, Adam, & Rigoux, 2014). The VBA toolbox tackles the problem of model fitting via a variational approximation that assumes factorized, Gaussian posteriors over the parameters (mean field/Laplace approximation), and provides the variational free energy as an approximation (lower bound) of the marginal likelihood. Our approach, instead, does not make any strong assumption, using MCMC to recover the full shape of the posterior, and state-of-the-art techniques to assess model performance.

Detailed, rigorous modeling of behavior is a necessary step to constrain the search for neural mechanisms implementing decision strategies (Krakauer, Ghazanfar, Gomez-Marin, MacIver, & Poeppel, 2017). We have provided a set of computational tools and demonstrated how they can be applied to answer specific questions about internal representation and decision strategies of the observer in multisensory perception, with the goal of increasing the set of models that can be investigated, and the robustness of such analyses. Thus, our tools can be of profound use not only to the field of multisensory perception, but to biological modeling in general.

## Methods

### Human psychophysics

#### Subjects

Eleven healthy adults (4 female; age 26.4 ± 4.6 years, mean ± SD) participated in the full study. Subjects had no previous history of neurological disorders and had normal or corrected-to-normal vision. Four other subjects completed only a partial version of the experiment, and their data were not analyzed here. The Institutional Review Board at the Baylor College of Medicine approved the experimental procedures and all subjects gave written informed consent.

#### Apparatus

Details of the experimental apparatus have been previously published and are only described here briefly (Dokka et al., 2015; Dokka, MacNeilage, DeAngelis, & Angelaki, 2011; Fetsch et al., 2009; MacNeilage, Zhang, DeAngelis, & Angelaki, 2012). Subjects were seated comfortably in a cockpit-style chair and were protectively restrained with a 5-point racing safety harness. Each subject wore a custom-made thermoplastic mesh mask that was attached to the back of the chair for head stabilization. The chair, a three-chip DLP projector (Galaxy 6; Barco) and a large projection screen (149 × 127 cm) were all mounted on a motion platform (6DOF2000E; Moog, Inc.). The projection screen was located ~ 65 cm in front of the eyes, subtending a visual angle of ~ 94° × 84°. Subjects wore LCD-based active 3D stereo shutter glasses (Crystal Eyes 4, RealD, Beverly Hills) to provide stereoscopic depth cues and headphones for providing trial timing-related feedback (a tone to indicate when a trial was about the begin and another when a button press was registered). This apparatus was capable of providing three self-motion conditions: vestibular (inertial motion through the movement of the platform), visual (optic flow simulating movement of the observer in a 3D virtual cloud of stars, platform stationary) and combined visual-vestibular heading (temporally-synchronized optic flow and platform motion) at various spatial discrepancies.

#### Stimuli

We modified a previous multi-sensory heading discrimination task (Fetsch et al., 2009). Here subjects experienced combined visual and vestibular translation in the horizontal plane (Figure 1A). The visual scene and platform movement followed a Gaussian velocity profile (displacement = 13 cm, peak Gaussian velocity = 26 cm/s and peak acceleration = 0.9m/s^2^, duration = 1 s). Visual and vestibular headings were either in the same direction or their movement trajectories were separated by a directional disparity, Δ, expressed in degrees (Figure 1A). The directional disparity Δ and visual cue reliability were varied on a trial-by-trial basis. Δ took one of five values, selected with equal probability: 0° (no conflict), 5°, 10°, 20° and 40°. Thus, visual and vestibular stimuli were in conflict in 80% of the trials. In each trial, Δ was randomly assigned to be positive (Figure 1A right, vestibular heading to the right of visual heading) or negative. Once a disparity value, Δ, was chosen, the mean heading angle 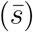, which represents the average of vestibular and visual headings, was uniformly randomly drawn from the discrete set {−25°, −20°,…, 25°}. Vestibular heading (*s*_vest,_ red trace in Figure 1) and visual heading (*s*_vis,_ black trace in Figure 1A) were generated by displacing the platform motion and optic flow on either side of the mean heading by Δ/2. The vestibular and visual headings experienced by subjects were defined as 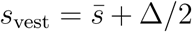 and 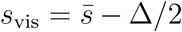, respectively. This procedure entailed that visual and vestibular heading directions presented in experiment were correlated (Figure 1B). Three levels of visual cue reliability (high, medium, and low) were tested. Visual reliability was manipulated by varying the percentage of stars in the optic flow that coherently moved in the specified heading direction. For all subjects, visual motion coherence at high reliability was set at 100%. For medium and low reliability, visual motion coherences ranged from 40-70% and 25-50%. Overall, there were 297 stimulus conditions (9 directional disparities × 11 mean heading directions × 3 visual cue reliabilities) which were randomly interleaved.

#### Tasks

First, subjects (*n* = 11) performed in a session of a unisensory heading discrimination task (left/right of straight ahead), in which visual or vestibular stimuli were presented in isolation. Vestibular stimuli had one fixed reliability level, whereas visual stimuli were tested on three different reliability levels, randomly interleaved, resulting in a total of 350 − 750 trials.

Then, subjects performed the explicit inference task (unity judgment). Here, subjects indicated if the visual and vestibular cues indicated heading in the same direction (“common” cause, *C* = 1) or in different directions (“different” causes, *C* = 2). Each combination of disparity and reliability was presented 30 times. Since each disparity was randomly assigned to be positive or negative on each trial, 0° disparity was presented 60 times at each visual cue reliability resulting in a total of 900 trials. Subjects did not receive feedback about the correctness of their responses.

Finally, the same subjects also participated in the implicit inference task – bisensory (inertial) discrimination. Here, subjects indicated the perceived direction of their inertial self-motion (left or right of straight ahead). Note that although both visual and vestibular stimuli were presented in each trial, subjects were asked to only indicate their perceived direction of inertial heading, similar to the bisensory auditory localization procedure in (Rohe & Noppeney, 2015b). Each combination of disparity and visual cue reliability was presented about 100 times. Since each disparity was randomly assigned to be positive or negative on each trial, 0° disparity was presented 200 times resulting in a total of about 3000 trials divided across several sessions. No feedback was given about the correctness of subjects’ responses.

#### Data analysis

For the explicit inference task, we computed the proportion of trials in which subjects perceived a common cause at each disparity and visual cue reliability. For the implicit inference task, we calculated the bias in perceived inertial heading. In order to compute the bias, we binned values of *s*_vis_ in the following intervals: {[−45°, −30°], [−27.5°, −22.5°], [−20°, −15°], [−12.5°, −7.5°], [−5°, −2.5°], 0°, [2.5°, 5°], [7.5°,12.5°], [15°, 20°], [22.5°, 27.5°], [30°, 45°]}. For each visual bin and level of visual cue reliability, we constructed psychometric functions by fitting the proportion of rightward responses as a function of *s*_vest_ with cumulative Gaussian functions (inset in Figure 3A). We defined the bias in the perceived inertial heading as minus the point of subjective equality (L/R PSE). A bias close to zero indicates that subjects accurately perceived their inertial (vestibular) heading. Substantial biases suggest that misleading visual cues exerted a significant influence on the accuracy of inertial heading discrimination. Repeated-measures ANOVA with disparity or visual bin and visual cue reliability as within-subjects factors were performed separately on the proportion of common cause reports and bias in perceived inertial heading. We applied Greenhouse-Geisser correction of the degrees of freedom in order to account for deviations from sphericity (Greenhouse & Geisser, 1959), and report effect sizes as partial eta squared, denoted with 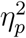. For all analyses the criterion for statistical significance was *p* < .05, and we report uncorrected *p*-values. Unless specified otherwise, summary statistics are reported in the text as mean ± SE between subjects.

### Causal inference models

We build upon standard causal inference (CI) models of multisensory perception (Körding et al., 2007). For concreteness, in the following description of CI models we refer to the visuo-vestibular example with binary responses (‘left/right’ for discrimination, and ‘yes/no’ for unity judgements). The basic component of any observer model is the trial response probability, that is the probability of observing a given response for a given trial condition (e.g., stimulus pair, uncertainty level, task). In the following we briefly review how these probabilities are computed.

All analysis code was written in MATLAB (Mathworks, Inc.), with core computations in C for increased performance (via *mex* files in MATLAB). Code is available at <u>https://github.com/lacerbi/visvest-causinf.</u>

#### Unisensory heading discrimination

We used subjects’ binary (‘left or right of straight forward’) heading choices, measured in the presence of visual-only and vestibular-only stimuli, to estimate subjects’ measurement noise in the respective sensory signals. Let us consider a trial with a vestibular-only stimulus (the computation for a visual-only stimulus is analogous). Subjects are asked whether the perceived direction of motion *s*_vest_ is to the left or to the right of straight forward (0°). We assume that the observer has access to a noisy measurement *x*_vest_ of stimulus *s*_vest_ (direction of motion), with probability density
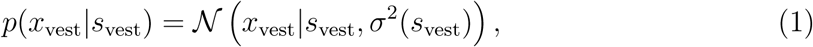

where 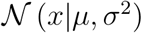 is a normal probability density with mean *μ* and variance *σ*^2^. Depending on the sensory noise model, the variance in Eq. 1 is either *constant* 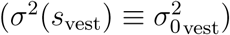 or *eccentricity-dependent* with base magnitude 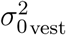 and noise that increases with eccentricity (distance from 0°) according to a parameter *w*_vest_ ≥ 0 (see Appendix A for details). For *w*_vest_ = 0, the eccentricity-dependent model reduces to the constant model. The observer’s posterior probability density over the vestibular stimulus is *p* (*s*_vest_|*x*_vest_) ∝ *p*(*x*_vest_|*s*_vest_)*p*_prior_(*s*_vest_), and we will see that under some assumptions the prior over heading directions is irrelevant for subsequent computations in the left/right unisensory task (see Appendix A).

We assume that observers compute the posterior probability that the stimulus is right of straight forward as 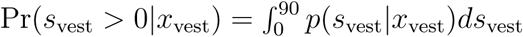, and respond ‘right’ if Pr(*s*_vest_ *>* 0|*x*_vest_) > 0.5; ‘left’ otherwise. Observers may also lapse and give a completely random response with probability λ (lapse rate). This yields
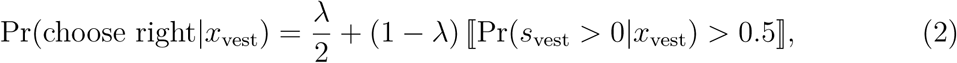

where 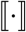 is Iverson bracket, which is 1 if the argument is true, and 0 otherwise (Knuth, 1992).

An analogous derivation is applied to each unisensory visual stimulus condition for respectively low, medium, and high visual reliability. We assume a distinct 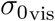 for each visual reliability condition, and, for the eccentricity-dependent models, a common *w*_vis_ for all visual reliability conditions, so as to reduce model complexity.

#### Unity judgment (explicit inference)

In a unity judgment trial, the observer explicitly evaluates whether there is a single cause (*C* = 1) underlying the noisy measurements *x*_vis_, *x*_vest_, or two separate causes (*C* = 2; see Figure 2B). All following probability densities are conditioned on *c*_vis_, the level of visual cue reliability in the trial, which is assumed to be known to the observer; we omit this dependence to reduce clutter. We consider three families of explicit CI strategies.

The *Bayesian* CI strategy computes the posterior probability of common cause
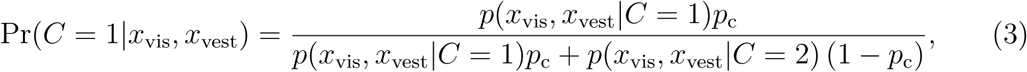

where 0 ≤ *p*_c_ ≡ Pr(*C* = 1) ≤ 1, the prior probability of a common cause, is a free parameter of the model. The derivation of *p*(*x*_vis_, *x*_vest_|*C* = *k*), for *k* = 1, 2, is available in Appendix A. The observer reports unity if the posterior probability of common cause is greater than 0.5, with the added possibility of random lapse,
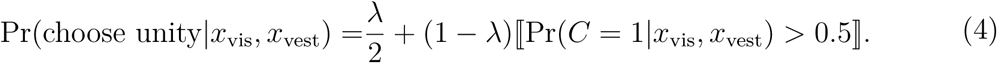

For a separate analysis we also considered a ‘probability matching’ variant that reports unity with probability equal to Pr(*C* = 1|*x*_vis_, *x*_vest_) (plus lapses).

As a non-Bayesian CI heuristic model, we consider a *fixed criterion* observer, who reports a common cause whenever the two noisy measurements are within a distance *κ*_c_ ≥ 0 from each other,

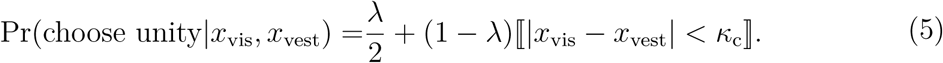

Crucially, the fixed criterion observer does not take into account stimulus reliability or other statistical information when inferring the causal structure.

Finally, we consider a *fusion* observer that eschews CI altogether. A classical ‘forced fusion’ observer would *always* report ‘unity’ in the explicit CI task, which is easily rejected by the data. Instead, we consider a *stochastic fusion* observer that reports ‘unity’ with probability *η*_low,_*η*_med,_ or *η*_high,_ depending only on the reliability of the visual cue, and discards any other information.

#### Bisensory inertial discrimination (implicit inference)

In bisensory inertial discrimination trials, the observer reports whether the perceived inertial heading *s*_vest_ is to the left or right of straight forward (0°). In this experiment, we do not ask subjects to report *s*_vis,_ but the inference would be analogous. The inertial discrimination task requires an implicit evaluation of whether there is a single cause to the noisy measurements *x*_vis_, *x*_vest_ (*C* = 1), or two separate causes (*C* = 2), for a known level of visual coherence *c*_vis_ (omitted from the notation for clarity).

If the observer knew that *C* = *k*, for *k* = 1, 2, the posterior probability density over the vestibular stimulus would be (see Appendix A)

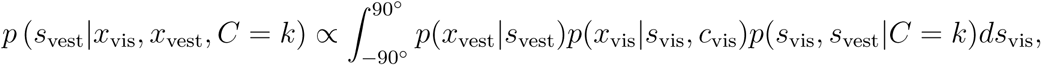

where the likelihoods are defined as per the uni-sensory task, Eq. 1, and for the prior over heading directions, *p*(*s*_vis_, *s*_vest_|*C*), see ‘Observers’ priors’ below.

The posterior probability of rightward motion is computed for *k* = 1, 2 as

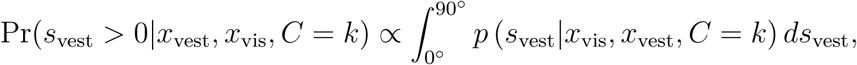

and an analogous equation holds for the posterior probability of leftward motion.

In general, the causal structure is implicitly inferred by the observer. We assume that observers combine cues according to
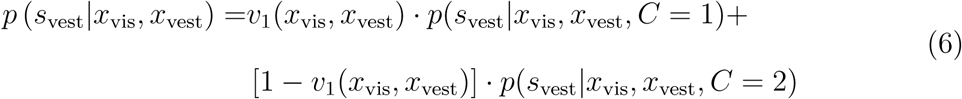

where 0 ≤ *v*_1_(*x*_vis_, *x*_vest_) ≤ 1 is the *implicit causal weight* associated by the observer to the hypothesis of a single cause, *C* = 1. The form of the causal weight depends on the observer’s implicit CI strategy.

We consider three families of implicit CI. For the *Bayesian* CI observer, the causal weight is equal to the posterior probability, *v*_1_(*x*_vis_, *x*_vest_) = Pr(*C* = 1|*x*_vis_, *x*_vest_), so that Eq. 6 becomes the expression for Bayesian model averaging (Körding et al., 2007; see Eq. 3 and Appendix A). As a variant of the Bayesian observer we consider a *probability matching* Bayesian strategy for which *v*_1_ = 1 with probability Pr(*C* = 1|*x*_vis_, *x*_vest_), and *v*_1_ = 0 otherwise. For the *fixed-criterion* observer, 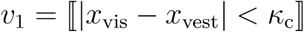, with *κ*_c_ ≥ 0 as per Eq. 5. Finally, for the *forced fusion* observer *v*_1_ ≡ 1.

The posterior probability of rightward motion is then 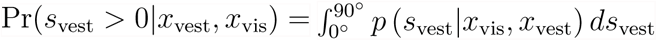, and an analogous equation holds for the posterior probability of leftward motion. We assume the observer reports the direction with highest posterior probability, with occasional lapses (see also Eq. 2),
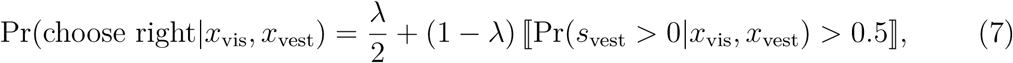

where *λ* ≥ 0 is the lapse rate.

#### Observers’ prior

We assume subjects develop a symmetric, unimodal prior over heading directions for unisensory trials. Due to the form of the decision rule (Eq. 2), a symmetric prior has no effect on the unisensory trials, so we only focus on the bisensory case.

For the bisensory prior over heading directions, *p*(*s*_vis_, *s*_vest_|*C*) we consider two families of priors. The *empirical* prior approximately follows the correlated structure of the discrete distribution of vestibular and visual headings presented in the experiment (Figure 1B). The *independent* prior assumes that observers learn a generic uncorrelated Gaussian prior over heading directions, as per Körding et al. (2007). See Appendix A for details.

#### Trial response probabilities

Eqs. 2, 4, 5, and 7 represent the probability that an observer chooses a specific response *r* (‘rightward’ or ‘leftward’ for discrimination trials, ‘same’ or ‘different’ for unity judgment trials), for given noisy measurements *x*_vis_ and *x*_v__est_ (or only one of the two for the unisensory task), and known visual reliability *c*_vis_. Since as experimenters we do not have access to subjects’ internal measurements, to compute the trial response probabilities we integrate (‘marginalize’) over the unseen noisy measurements for given heading directions *s*_vis_ and *s*_vest_ presented in the trial.

For the unisensory case, considering as example the vestibular case, we get
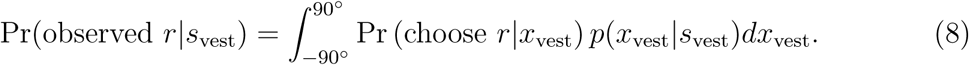

For the bisensory case, either unity judgment or inertial discrimination, we have
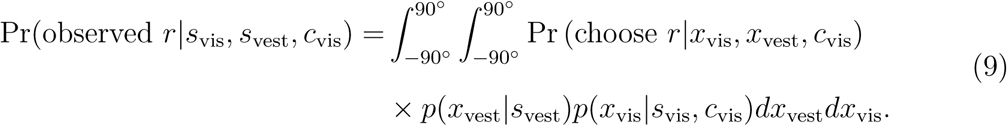

It is customary in the causal inference literature to approximate these integrals via Monte Carlo sampling, by drawing a large number of noisy measurements from the noise distributions (e.g., de Winkel et al., 2015; Körding et al., 2007; Rohe & Noppeney, 2015a; Wozny et al., 2010). Instead, we computed the integrals via numerical integration, which is more efficient than Monte Carlo techniques for low dimensional problems (Press, Flannery, Teukolsky, & Vetterling, 2007).

We used the same numerical approach to evaluate Eqs. 2, 4, 5, and 7, including an adaptive method for choice of integration grid. All numerical integrals were then coded in C (*mex* files in MATLAB) for additional speed. See Appendix B for computational details.

### Model fitting

For a given model, we denote its set of parameters by a vector ***θ.*** For a given model and dataset, we define the parameter log likelihood function as
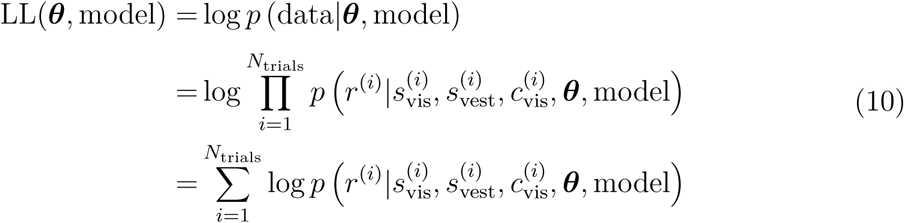

where we assumed conditional independence between trials; *r*^(*i*)^ denotes the subject’s response (‘right’ or ‘left’ for the discrimination trials; ‘common’ or ‘separate’ causes in unity judgment trials); 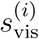 and 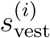 are, respectively, the direction of motion of the visual (resp. vestibular) stimulus (if present), and 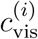 is the visual coherence level (that is, reliability: low, medium, or high), in the *i*-th trial.

#### Maximum likelihood estimation

First, we fitted our models to the data via maximum likelihood estimation (MLE), by finding the parameter vector ***θ\**** that maximizes the log likelihood in Eq. 10. For optimization of the log likelihood, we used Bayesian Adaptive Direct Search (BADS; <u>https://github.com/lacerbi/bads;</u> Acerbi & Ma, 2017). BADS is a black-box optimization algorithm that combines a mesh-adaptive direct search strategy (Audet & Dennis Jr, 2006) with a local Bayesian optimization search step based on Gaussian process surrogates (see Brochu, Cora, & De Freitas, 2010; Shahriari et al., 2016 for an introduction to Bayesian optimization). Bayesian optimization is particularly useful when the target function is costly to evaluate or the likelihood landscape is rough, as it is less likely to get stuck in local optima than other algorithms, and may reduce the number of function evaluations to find the (possibly global) optimum. In our case, evaluation of the log likelihood function for a single parameter vector ***θ*** could take up to ~ 2-3 s for bisensory datasets, which makes it a good target for BO. We demonstrated in a separate benchmark that BADS is more effective than a large number of other MATLAB optimizers for our problem (‘causal inference’ problem set in Acerbi & Ma, 2017). See Appendix B for more details about the algorithm and the optimization procedure.

For each subject we first fitted separately the datasets corresponding to three tasks (unisensory and bisensory heading discrimination, unity judgment), and then performed joint fits by combining datasets from all tasks (summing the respective log likelihoods).

#### Posterior sampling

As a complementary approach to ML parameter estimation, for each dataset and model we calculated the posterior distribution of the parameters,
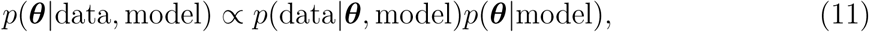

where *p*(data|***θ***, model) is the likelihood (see Eq. 10) and *p*(***θ***|model) is the prior over parameters. We assumed a factorized prior 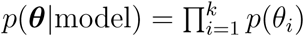 and a non-informative uniform prior over a bounded interval for each model parameter (uniform in log space for scale parameters such as all noise base magnitudes, fixed criterion *κ*_c_, and prior parameters *σ*_prior_ and Δ_prior_); see Table 1.

We approximated Eq. 11 via Markov Chain Monte Carlo (MCMC) sampling. We used a custom-written sampling algorithm that combines slice sampling (Neal, 2003) with adaptive direction sampling (ADS; Gilks et al., 1994) and a number of other tricks (https://github.com/lacerbi/eissample). Slice sampling is a flexible MCMC method that, in contrast with the common Metropolis-Hastings transition operator, requires very little tuning in the choice of length scale. ADS is an ensemble MCMC method that shares information between several dependent chains (also called ‘walkers’; Foreman-Mackey, Hogg, Lang, & Goodman, 2013) in order to speed up mixing and exploration of the state space. For details about the MCMC algorithm and the sampling procedure, see Appendix B.

### Factorial model comparison

We built different observer models by factorially combining three factors: CI strategy (Bayesian, fixed-criterion, or fusion); shape of sensory noise (constant or eccentricity-dependent); and type of prior over heading directions (empirical or independent); see Figure 2A and ‘Causal inference models’ section of the Methods for a description of the different factors.

For each subject, we fitted the different observer models, first separately to different tasks (unity judgment and bisensory inertial discrimination), and then performed a joint fit by combining datasets from all tasks (including the unisensory discrimination task). We evaluated the fits with a number of model comparison metrics and via an objective goodness of fit metric. Finally, we combined evidence for different model factors across subjects with a hierarchical Bayesian approach.

We verified our ability to distinguish different models with a model recovery analysis, described in Appendix B.

#### Model comparison metrics

For each dataset and model we computed a number of different model comparison metrics, all of which take into account quality of fit and penalize model flexibility, but with different underlying assumptions.

Based on the maximum likelihood solution, we computed Akaike information criterion with a correction for sample size (AICc) and Schwarz’s ‘Bayesian’ Information criterion (BIC),
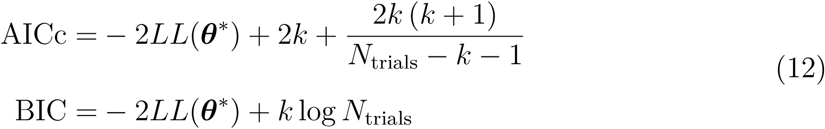

where *N*_trials_ is the number of trials in the dataset and *k* is the number of parameters of the model. The factor of −2 that appears in both definitions is due to historical reasons, so that both metrics have the same scale of the deviance.

To assess model performance on unseen data, we performed Bayesian leave-one-out (LOO) cross-validation. Bayesian LOO cross-validation computes the posterior of the parameters given *N*_trials_ − 1 trials (training), and evaluates the (log) expected likelihood of the left-out trial (test); the procedure is repeated for each trial, yielding the leave-one-out score
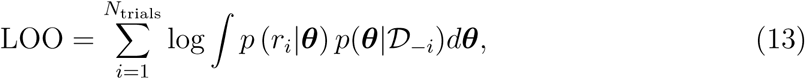

where *p*(*r*_*i*_|***θ***) is the likelihood associated to the *i*-th trial alone, and 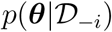 is the posterior over ***θ*** given all trials except the *i*-th one. Eq. 13 can be estimated at prohibitive computational cost by separately sampling from the leave-one-out posteriors via *N*_trials_ distinct MCMC runs. A more feasible approach comes from noting that all posteriors differ from the full posterior by only one data point. Therefore, the leave-one-out posteriors can be approximated via *importance sampling*, reweighting the full posterior obtained via MCMC. However, a direct approach of importance sampling can be unstable, since the full posterior is typically narrower than the leave-one-out posteriors. Pareto-smoothed importance sampling (PSIS) is a recent technique to stabilize the importance weights (Vehtari et al., 2015), implemented in the psisloo package (https://github.com/avehtari/PSIS). Thus, Eq. 13 is approximated as
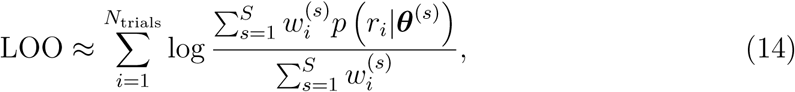

where ***θ***^(*s*)^ is the *s*-th parameter sample from the posterior, and 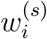 are the Pareto-smoothed importance weights associated to the *i*-th trial and *s*-th sample (out of *S*); see Vehtari et al. (2016) for details. PSIS also returns for each trial the exponent *k_i_* of the fitted Pareto distribution; if *k_i_* is greater than 1 the moments of the importance ratios distribution do not exist and the variance of the PSIS estimate is finite but may be large; this provides a natural diagnostic for the method (Vehtari et al., 2016; see Appendix B). LOO is our comparison metric of choice (see Discussion). LOO scores for all models and subjects are reported in Appendix D.

Finally, we approximated the *marginal likelihood* of the model,
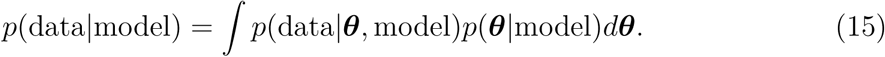

The marginal likelihood is a common metric of model evidence that naturally incorporates a penalty for model complexity due to Bayesian Occam razor (MacKay, 2003). However, the integral in Eq. 15 is notoriously hard to evaluate. Here we computed an approximation of the log marginal likelihood (LML) based on MCMC samples from the posterior, by using a *weighted* harmonic mean estimator (Robert et al., 2009). The formula for the approximation is
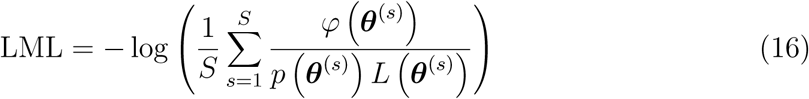

where the sum is over *S* samples from the posterior, ***θ***^(*s*)^ is the *s*-th sample, *p*(***θ***) the prior, *L*(***θ***) the likelihood, and *φ*(***θ***) is an arbitrary weight probability density. The behavior of the approximation depends crucially on the choice of *φ*; it is important that *φ* has thinner tails than the posterior, lest the variance of the estimator grows unboundedly. We followed the suggestion of Robert et al. (2009) and adopted a finite support distribution over a high posterior density (HPD) region. We fitted a variational Gaussian mixture model to the posterior samples (Bishop, 2006; <u>https://github.com/lacerbi/vbgmm),</u> and then we replaced each Gaussian component with a uniform distribution over an ellipsoid region proportional to the covariance matrix of the component. The proportionality constant, common to all components, was picked by minimizing the empirical variance of the sum in Eq. 16 (Caldwell & Liu, 2014).

#### Hierarchical Bayesian model selection

We performed Bayesian model selection (BMS) at the group level via a hierarchical approach that treats subjects and models as random variables (Stephan et al., 2009). BMS infers the posterior over model frequencies in the population, expressed as Dirichlet distributions parametrized by the concentration parameter vector ***α.*** As a summary statistic we consider the protected exceedance probability 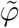, that is the probabilty that a given model or model factor is the most likely model or model factor, above and beyond chance (Rigoux et al., 2014). For the *i*-th model or model factor,

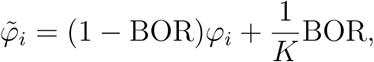

where *K* is the number of models (or model factors), *φ_i_* is the unprotected exceedance probability for the *i*-th model or model factor (Stephan et al., 2009), and BOR is the Bayesian omnibus risk – the posterior probability that the data may be explained by the null hypothesis according to which all models (or model factors) have equal probability (Rigoux et al., 2014). For completeness, we report posterior model frequencies and BOR in the figures, but we do not focus on model frequencies per se since our sample size does not afford a more detailed population analysis.

To compute the posterior over model factors in the population we exploit the agglomerative propery of the Dirichlet distribution, and sum the concentration parameters of models that belong to the same factor component (Stephan et al., 2009). While the agglomerative property allows to easily compute the posterior frequencies and the *unprotected* exceedance probabilities for each model factor, calculation of the protected exceedance probabilities required us to compute the BOR for the model factor setup (the probability that the observed differences in factor frequencies may have arisen due to chance).

Additionally, the BMS method requires to specify a Dirichlet prior over model frequencies, represented by a concentration parameter vector *α*_0_ · ***w***, with *w_k_* = 1 for any model *k* and *α*_0_ > 0. The common choice is *α*_0_ = 1 (flat prior over model frequencies), but given the nature of our factorial analysis we prefer a flat prior over model factors (*α*_0_ = average number factors / number of models), where the average number of factors is ≈ 2.33 for the bisensory tasks and ≈ 2.67 for the joint fits. This choice entails that the concentration parameter of the agglomerate Dirichlet distributions, obtained by grouping models that belong to the same factor component, is of order ~ 1 (it cannot be exactly one since different factors have different number of components). When factor components within the same factor had unequal numbers of models, we modified the prior weight vector ***w*** such that every component had equal prior weight. We verified that our main results did not depend on the specific choice of Dirichlet prior (Figure 7, third row).

#### Parameter compatibility metric

Before performing the joint fits, we tested whether model parameters differed across the three tasks (unisensory and bisensory discrimination, unity judgment). On one end of the spectrum, the fully Bayesian approach would consist of comparing all combinations of models in which parameters are shared vs. distinct across tasks, and check which combination best explains the data. However, this approach is intractable in practice due to the combinatorial explosion of models, and undesirable in theory due to the risk model overfitting. On the simplest end of the spectrum, we could look at the credible intervals of the parameter posteriors for each subject and visually check whether they are mostly overlapping for different tasks.

As a middle ground, we computed separately for each parameter what we defined as the *compatibility probability C_p_*, that is the probability that for most subjects the parameter is exactly the same across tasks (*H*_0_), as opposed to being different (*H*_1_), above and beyond chance.

For a given subject, let *y*_1_, *y*_2_, and *y*_3_ be the datasets of the three tasks. For a given parameter *θ* (e.g., lapse rate), we computed the compatibility likelihoods
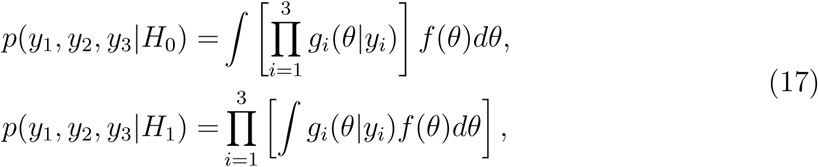

where *g_i_*(***θ****|y_i_*) is the marginal posterior over ***θ*** for the dataset *y_i_*, and *f*(***θ***) is the prior over ***θ***. Having computed the compatibility likelihoods for all subjects, we defined *C_p_* as the protected exceedance probability of model *H*_0_ vs. model *H*_1_ for the entire group.

For each subject and task, the marginal posteriors *g_i_*(***θ***| *y_i_*) were obtained as a weighted average over models, with weight equal to each model’s posterior probability for that subject according to the group BMS method via LOO, and considering only the subset of models that include the parameter of interest (see Figure 5).

For the prior *f*(***θ***) over a given parameter ***θ***, for the purposes of this analysis only, we followed an empirical Bayes approach informed by the data and use a truncated Cauchy prior fitted to the average marginal posterior of ***θ*** across subjects, defined over the range of the MCMC samples for ***θ***.

#### Absolute goodness of fit

Model comparison yields only a *relative* measure of goodness of fit, but does not convey any information of whether a model is a good description of the data in an absolute sense. A standard metric such as the coefficient of variation *R*^2^ is not appropriate for binary data. Instead, we extended the approach of Shen and Ma (2016) and defined *absolute goodness of fit* as
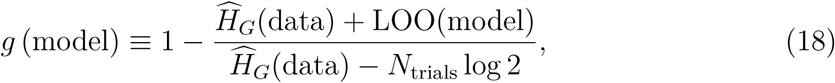

where 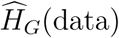 is an estimate of the entropy of the data obtained via Grassberger’s estimator (Grassberger, 2003) and LOO(model) is the LOO score of the model of interest.

The numerator in Eq. 18 represents the Kullback-Leibler (KL) divergence between the distribution of the data and the distribution predicted by the model (that is, how well the model captures the data), which is compared as a reference to the KL divergence between the data and a chance model (at the denominator). See Appendix C for a derivation of Eq. 18, and code is available at <u>https://github.com/lacerbi/gofit.</u>

### The cookbook

The Bayesian cookbook for causal inference (CI) in multisensory perception, or simply ‘the cookbook’, consists of a number of algorithms and computational techniques to perform efficient and robust Bayesian comparison of CI models. We applied and demonstrated these methods at different point in the main text; further details can be found here in the Methods and Appendices. For reference, we summarize the main techniques of interest in Table 2.

**Table 2.** List of algorithms and computational procedures. List of useful algorithms and computational procedures.

| Description | Code | References |
| --- | --- | --- |
| *Model fitting* |  |  |
| Efficient computation of log likelihood | https://github.com/lacerbi/visvest-causinf | This work |
| Maximum-likelihood estimation (optimization) | https://github.com/lacerbi/bads | Acerbi and Ma (2017) |
| Posterior estimation (MCMC sampling) | https://github.com/lacerbi/eissample | In preparation |
| *Model evaluation and comparison* |  |  |
| Leave-one-out cross validation (LOO) | https://github.com/avehtari/PSIS | Vehtari et al. (2015, 2016) |
| Estimate of the marginal likelihood | https://github.com/lacerbi/marglike | Robert et al. (2009), in preparation |
| Parameter compatibility test | https://github.com/lacerbi/comprob | This work |
| Objective goodness of fit | https://github.com/lacerbi/gofit | Shen and Ma (2016), this work |
| Group Bayesian Model Selection (BMS) | *spm_BMS* function in the SPM12 package<br>http://www.fil.ion.ucl.ac.uk/spm/ | Stephan et al. (2009),<br>Rigoux et al. (2014) |

## Appendix A Observer model factors

In this appendix we describe details of the factors used to build observer models.

### Sensory noise

For a given modality mod ∊ {vis, vest}, the measurement noise distribution has the shape
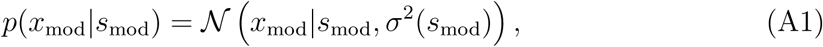

where 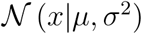 is a normal distribution with mean *μ* and variance *σ*^2^. Note that for a visual stimulus the measurement distribution and the variance in Eq. A1 also depend on the visual coherence level *c*_vis_ in the trial, such that *σ*^2^(*s*_vis_) ≡ *σ*^2^(*s*_vis_*, c*_vis_), but in the following we omit this dependence to simplify the notation.

For the variance we consider two possible models,
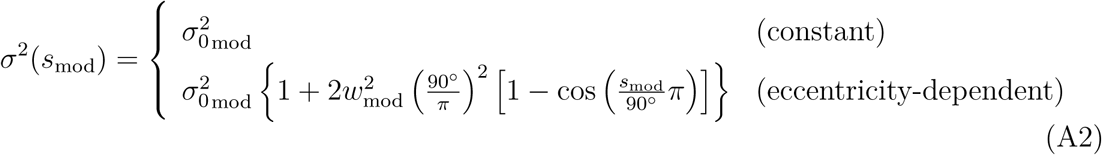

where 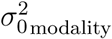 is the base variance and *w*_mod_ is related to the Weber fraction near 0°. In fact, for small values of *s*_mod,_ Eq. A2 reduces to 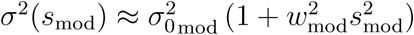, which is a generalized Weber’s law.

The broad shape of the chosen periodic formula for the eccentriticy-dependent noise model, which peaks at ±90°, derives from empirical results in a visuo-vestibular task with the same apparatus with human and monkey subjects (see Figure 2 in Gu, Fetsch, Adeyemo, DeAngelis, & Angelaki, 2010; see also Crane, 2012). We note that our noise shape differs from that adopted in other works (with different setups), which used a sinusoidal with twice the frequency that peaks at ±45°, ±135’ (de Winkel et al., 2015, 2017). Since in our setup the heading directions were restricted to the ±45° range (with most directions in the ±25° range), the exact shape of periodicity is largely irrelevant, but understanding differences in noise models may be important for experiments with wider heading direction ranges. Moreover, due to the limited range of directions in our experiment, we can afford to use Gaussian distributions, which allow for faster and simpler computations, as opposed to von Mises (circular normal) distributions which would be otherwise more appropriate for a fully circular domain.

All constant noise models have four parameters (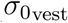, and a separate 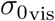 for each visual coherence level, low, medium and high). Eccentricity-dependent models have two additional parameters, *w*_vest_ and *w*_vis_ (the latter is common to all visual stimuli, to prevent overfitting).

### Prior

For unisensory trials, we assume that observers have a unimodal symmetric prior over heading directions, peaked at 0° (the exact shape is irrelevant). Due to the form of the decision rule, such prior has no influence over the unisensory left/right discrimination task.

For bisensory trials (both unity judgment and inertial discrimination tasks), we consider two alternative models for priors. The *empirical* prior consists of an approximation of the actual prior used in the experiment, that is
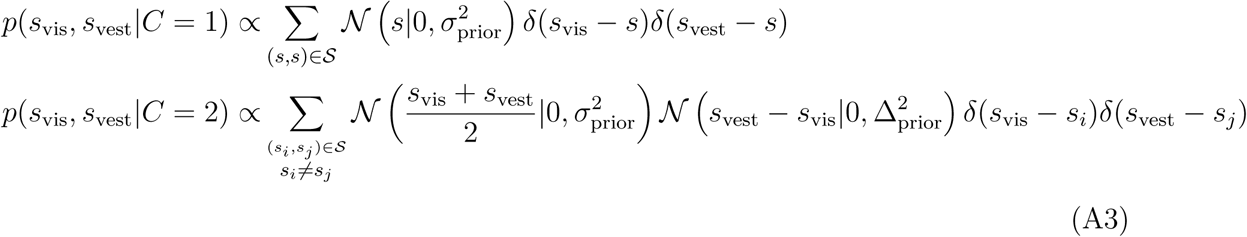

where 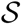 is the discrete set of pairs of visual and vestibular headings in the experiment. The two equations consider respectively only diagonal elements (equal heading directions, *C* = 1) or off-diagonal elements (different directions, *C* = 2) of Figure 1B in the main text. The approximation here is given by the two Gaussian distributions (defined on the discrete set), which impose additional shrinkage for the mean of the stimuli (governed by 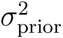 and for the disparity (governed by 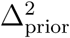. For 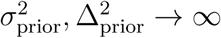, Eq. A3 converges to the distributions of directions used in the experiment for *C* = 1 and *C* = 2.

Alternatively, we consider an *independent* prior, that is
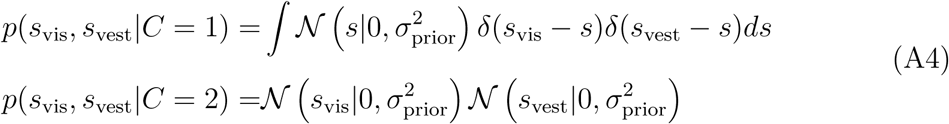

which assumes observers build a single prior over heading directions which is applied independently to both modalities (Körding et al., 2007). The first integral is a formal way to impose *s* ≡ *s*_vis_ = *s*_vest_.

We note that a continuous approximation of Eq. A3 may seem more realistic than the adopted discrete distribution of directions. However, an observer model with a correlated, continuous prior is computationally intractable since evaluation of the log likelihood involves a non-analytical four-dimensional integral, which increases the computational burden by an order of magnitude. As a sanity check, we implemented observers that use a continuous approximation of Eq. A3 and verified on a subset of observers and models that results of model fits and model predictions were indeed nearly identical to the discrete case.

Independent prior models have one parameter *σ*_prior_ for the width of the prior over headings. Empirical prior models have an additional parameter Δ_prior_ for the width of the prior over disparities.

### Causal inference strategy

The basic causal inference strategies: *Bayesian, fixed-criterion* and *fusion* are described in the main text. We report here some additional definitions and derivations. All integrals in this section are in the [−90°, 90°] range, unless noted otherwise.

#### Posterior probability of causal structure

For a Bayesian observer, the posterior probability of common cause is

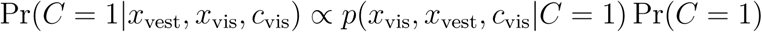

where Pr(*C* = 1) ≡ *p*_c_, the prior probability of a common cause, is a free parameter of the model. Then
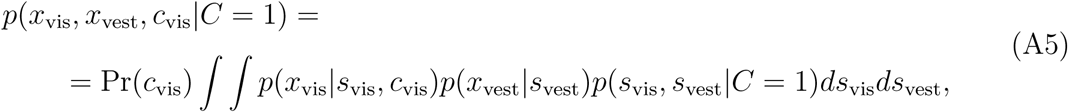

where the likelihoods are defined by Eq. A1, the prior is defined by Eqs. A3 and A4, and 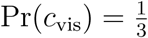. For the independent prior case we can further simplify

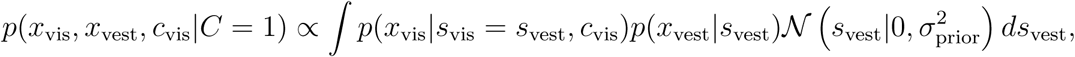

whereas the solution for the empirical prior is similar, but with a sum over the discrete stimuli such that *s*_vis_ = *s*_vest_.

Conversely, the
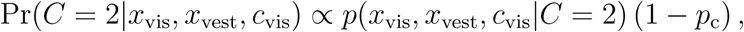

Where
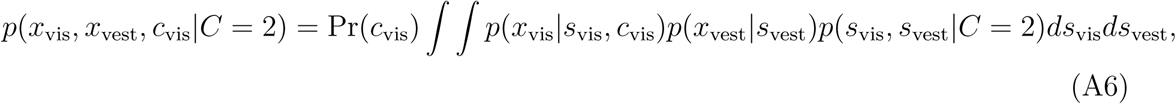

which for the independent prior becomes

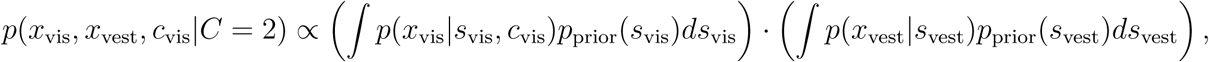

that is the product of two one-dimensional integrals. For the empirical prior Eq. A6 does not simplify, but becomes a discrete sum over 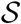 (see Eq. A3).

#### Posterior probability of left/right discrimination (*C* = 1)

In bisensory inertial discrimination trials the observer may implicitly contemplate two scenarios: that there is only one common cause (*C* = 1), or that there are two distinct causes (*C* = 2). We consider inference in the two separate scenarios, and then see how the observer can combine them.

For *C* = 1, the observer’s posterior probability density over over the inertial heading direction is
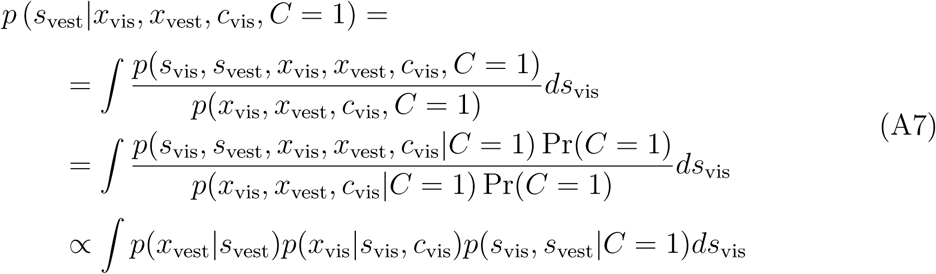

which for the independent prior becomes

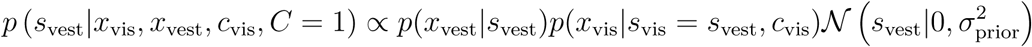

and the solution is similar for the empirical prior, constraining *s*_vest_ to take only the discrete values used in the experiment for *C* = 1.

#### Posterior probability of left/right discrimination (*C* = 2)

For *C* = 2, the observer’s posterior over inertial heading is
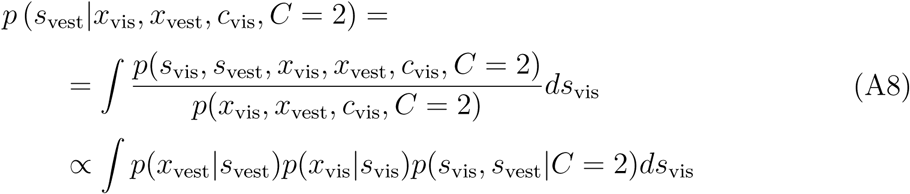

which for the independent prior can be further simplified as

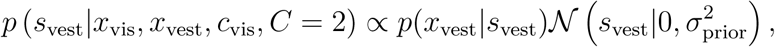

whereas for the empirical prior the integral in Eq. A8 becomes a sum over discrete pairs of heading directions used in the experiment.

#### Posterior probability of left/right discrimination (*C* unknown)

If the causal structure is unknown, a Bayesian observer that follows a ‘model averaging’ strategy marginalizes over possible causal structures (here, *C* = 1 and *C* = 2; Körding et al., 2007. The observer’s posterior probability density over the inertial heading direction is
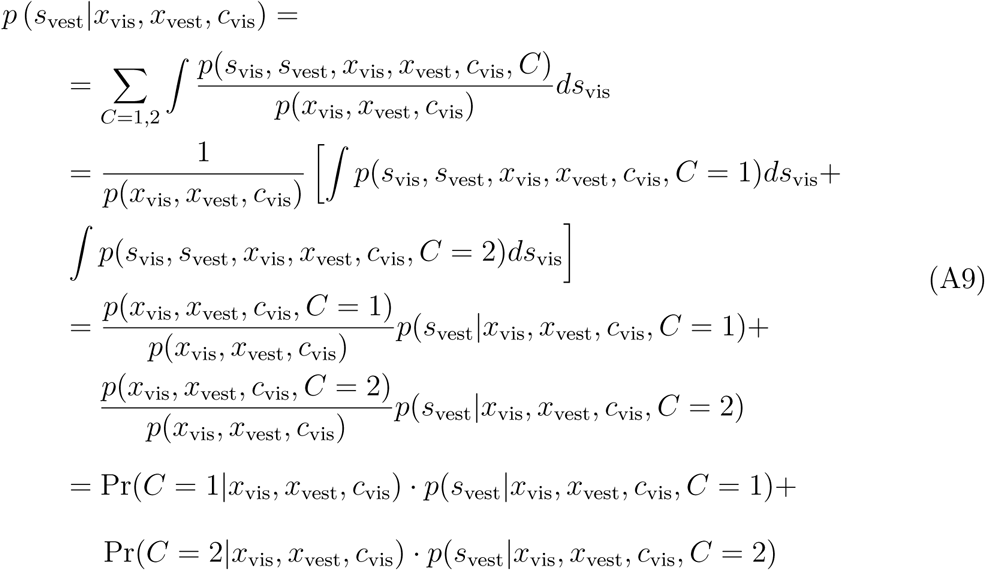

where *p*(*s*_vest_|*x*_vis_, *x*_vest_, *c*_vis_, *C*) has been defined in the previous subsections and Pr(*C*|*x*_vis_, *x*_vest_, *c*_vis_) is the posterior over causal structures.

We generalize Eq. A9 as

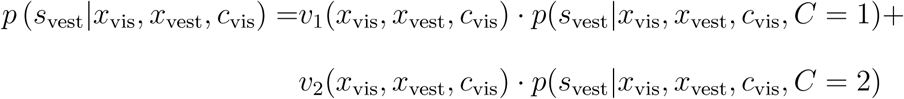

where *v_k_*(*x*_vis_, *x*_vest_, *c*_vis_), for *k* = 1, 2, are the posterior *causal weights* assigned by the observer to the two causal structures, with *v*_2_(*x*_vis_, *x*_vest_, *c*_vis_) = 1 − *v*_1_(*x*_vis_, *x*_vest_) and 0 ≤ *v*_1_ (*x*_vis_, *x*_vest_, *c*_vis_) ≤ 1. For a Bayesian observer, the causal weights are equal to the posterior probabilities (Eq. A9); in the main text we describe other models.

### Model parameters

All models except stochastic fusion have five parameters ***θ***_default_ by default: three visual base noise parameters 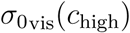, 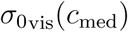, and 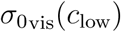; a vestibular base noise parameter 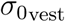; and a lapse rate λ.

| Observer model | Parameters | # |
| --- | --- | --- |
| Bayesian (*unisensory only*) | $\theta_{\text{default}}$ | 5 |
| Bayesian CI | $\theta_{\text{default}}, \sigma_{\text{prior}}, p_{\text{c}}$ | 7 |
| Fixed-criterion CI | $\theta_{\text{default}}, \kappa_{\text{c}}$ | 6 |
| Fusion CI | $\theta_{\text{default}}$ | 5 |
| Stochastic fusion (*unity judgment only*) | $\eta_{\text{high}}, \eta_{\text{med}}, \eta_{\text{low}}$ | 3 |
| *Add-ons* |  |  |
| with eccentricity-dependent noise | $+\{w_{\text{vis}}, w_{\text{vest}}\}$ | +2 |
| with empirical priors (*Bayesian*) | $+\{\Delta_{\text{prior}}\}$ | +1 |
| with empirical priors (*non-Bayesian*) | $+\{\sigma_{\text{prior}}, \Delta_{\text{prior}}\}$ | +2 |

## Appendix B Computational details

We describe in this appendix a number of computational and algorithmic details.

### Integrals

Due to lack of analytical solutions, we computed all one-dimensional and two-dimensional integrals numerically, via either Simpson’s or trapezoidal rule with a equi-spaced grid on the integration domain (Press et al., 2007). We had two types of integrals: integrals over *x*_vis_, *x*_vest_ for marginalization over the noisy stimuli, and integrals over *s*_vis_ and/or *s*_vest_ for computation of the observer’s decision rule (Eqs. A5, A6, A7 and A8).

For marginalization over noisy measurement *x*_vis_ and *x*_vest,_ we used a regular 401 × 401 grid for which we adjusted the range of integration in each modality to up to 5 SD from the mean of the noisy measurement distribution (or ±180°, whichever was smaller). For large noise, we used wrapped normal distributions.

For computation of the decision rule, we assumed that observers believed, due to the experimental setup and task instructions, that the movement direction would be forward, so limited to the ±90° range. We adjusted the integration grid spacing Δ*s* (hence the number of grid points) adaptively for each parameter vector ***θ***, defining

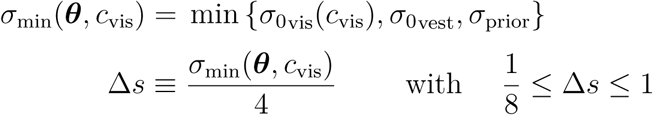

and we rounded Δ*s* to the lowest exact fraction of the form 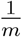, with 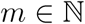 and 1 ≤ *m* ≤ 8. The above heuristic afforded fast and accurate evaluation of the integrals, since the grid spacing was calibrated to be smaller than the length scale of the involved distributions (measurement noise and prior).

Finally, we note that we tried other standard numerical integration methods which were ineffective. Gauss-Hermite quadrature (Press et al., 2007) led to large numerical errors because the integrand is discontinuous and bounded, a very bad fit for a polynomial. Global adaptive quadrature methods (such as quad in MATLAB, and other custom-made implementations) were simply too slow, even when reducing the requested precision. We coded all two-dimensional numerical integrals in C (via *mex* files in MATLAB) for maximal performance.

### Optimization

For optimization of the log likelihood (maximum-likelihood estimation, or MLE), we used Bayesian Adaptive Direct Search (BADS; <u>https://github.com/lacerbi/bads;</u> Acerbi & Ma, 2017). BADS follows a mesh adaptive direct search (MADS) procedure that alternates poll steps and search steps. In the poll step, points are evaluated on a (random) mesh by taking one step in one coordinate direction at a time, until an improvement is found or all directions have been tried. The step size is doubled in case of success, halved otherwise. In the search step, a Gaussian process (GP) is fit to a (local) subset of the points evaluated so far. Points to evaluate during the search are iteratively chosen by maximizing the predicted improvement (with respect to the current optimum) over a set of candidate points. Adherence to the MADS framework guarrantees convergence to a (local) stationary point of a noiseless function under general conditions (Audet & Dennis Jr, 2006). The basic scheme is enhanced with heuristics to accelerate the poll step, to update the GP hyper-parameters, to generate a good set of candidate points in the search step, and to deal robustly with noisy functions. See Acerbi and Ma (2017) for details.

For each optimization run, we initialized our algorithm by randomly choosing a point inside a hypercube of plausible parameter values in parameter space. We refined the output of each BADS run with a run of patternsearch (MATLAB). To avoid local optima, for each optimization problem we performed 150 independent restarts of the whole procedure and picked the highest log likelihood value.

As a heuristic diagnostic of global convergence, we computed by bootstrap the value of the global optimum we would have found had we only used *n*_r_ restarts, with 1 ≤ *n*_r_ ≤ 150. We define the ‘estimated regret’ as the difference between the actual best value of the log likelihood found and the bootstrapped optimum. For each optimization problem, we computed the minimum value 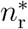 for which the probability of having an estimated regret less than 1 was 99% (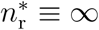 if such *n*_r_ does not exist). The rationale is that if the optimization landscape presents a large number of local optima, and new substantially improved optima keep being found with increasing *n*_r_, the bootstrapped estimated regret would keep changing with *n*_r_, and 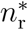 would be 150 or ∞. For almost all optimization problems, we found 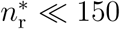. This suggests that the number of restarts was large enough; although no optimization procedure in a non-convex setting can guarantee convergence to a global optimum in a finite time without further assumptions.

### Markov Chain Monte Carlo (MCMC) sampling

As a complementary approach to MLE model fitting, for each dataset and model we calculated the posterior distribution of the parameters via MCMC (see main text).

We used a custom-written sampling algorithm that combines slice sampling (Neal, 2003) with adaptive direction sampling (ADS; Gilks et al., 1994).^1^ Slice sampling is a flexible MCMC method that, in contrast with the common Metropolis-Hastings transition operator, requires very little tuning in the choice of length scale. ADS is an ensemble MCMC method that shares information between several dependent chains (also called ‘walkers’; Foreman-Mackey et al., 2013) in order to speed up mixing and exploration of the state space. For each ensemble we used 2(*p* + 1) walkers, where *p* is the number of parameters of the model. Walkers were initialized to a neighborhood of the best local optima found by the optimization algorithm. Each ensamble was run for 10^4^ to 2.5 · 10^4^ burn-in steps that were discarded, after which we collected 5 · 10^3^ to 10^4^ samples per ensemble.

At each step, our method iteratively selects one walker in the ensemble and first attempts an independent Metropolis update. The proposal distribution for the independent Metropolis is a variational mixture of Gaussians (Bishop, 2006) fitted to a fraction of the samples obtained during burn-in via the `vbgmm` toolbox for MATLAB.^2^ Note that the proposal distribution is fixed at the end of burn-in and does not change thereafter, ensuring that the Markov property is not affected (although non-Markovian adaptive MCMC methods could be applied; see Andrieu & Thoms, 2008). After the Metropolis step, the method randomly applies with probability 1/3 one of three Markov transition operators to the active walker: coordinate-wise slice sampling (Neal, 2003), parallel-direction slice sampling (MacKay, 2003), and adaptive-direction slice sampling (Gilks et al., 1994; Neal, 2003). We also fit a variational Gaussian mixture model to the last third of the samples at the end of the burnin period, and we used the variational mixture as a proposal distribution for an independent Metropolis step which was attempted at every step.

For each dataset and model, we ran three independent ensembles. We visually checked for convergence the marginal pdfs and distribution of log likelihoods of the three sampled chains. For all parameters, we computed Gelman and Rubin’s potential scale reduction statistic *R* and effective sample size *n*_eff_ (Gelman et al., 2013) using Simo Särkkä and Aki Vehtari’s psrf function for MATLAB.^3^ For each dataset and model, we looked at the largest *R*(*R*_max_) and smallest 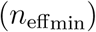 across parameters. Large values of *R* indicate convergence problems whereas values close to 1 suggest convergence. *n*_eff_ is an estimate of the actual number of independent samples in the chains; a few hundred independent samples are sufficient for a coarse approximation of the posterior (Gelman et al., 2013). Longer chains were run when suspicion of a convergence problem arose from any of these methods. Samples from independent ensembles were then combined (thinned, if necessary), yielding 1.5 · 10^4^ posterior samples per dataset and model. In the end, average *R*_max_ (across datasets and models) was ~ 1.002 (range: [1.000 − 1.035]), suggesting good convergence. Average 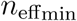 was ~ 8881 (range: [483 − 15059]), suggesting that we had obtained a reasonable approximation of the posteriors.

### Pareto smoothed importance sampling diagnostics

As our main metric of model comparison we computed the Bayesian leave-one-out cross-validation score (LOO) via Pareto-smoothed importance sampling (PSIS; Vehtari et al., 2015, 2016); see Methods in the main text.

For a given trial 1 ≤ *i* ≤ *N*_trials_, with *N*_trials_ the total number of trials, the PSIS approximation may fail if the leave-one-out posterior differs too much from the full posterior. As a natural diagnostic, PSIS also returns for each trial the exponent *k_i_* of the fitted Pareto distribution. If *k_i_* > 0.5 the variance of the raw importance ratios distribution does not exist, and for *k_i_* > 1 also the mean does not exist. In the latter case, the variance of the PSIS estimate is still finite but may be large. In practice, Vehtari et al. suggest to double-check trials with *k_i_* > 0.7 (Vehtari et al., 2016).

Across all our models and datasets, we found 2382 trials out of 1137100 with *k_i_* > 0.7 (0.21%). We examined the problematic trials, finding that the issue was in almost all cases the discontinuity of the observer’s decision rule. For all problematic trials the LOO*_i_* scores were compatible with the values found for non-problematic trials, suggesting that the variance of the PSIS estimate was still within an acceptable range. We verified on a subset of subjects that the introduction a softmax with small spatial constant on the decision rule would remove the discontinuity and the problems with Pareto fitting, without significantly affecting the LOO*_i_* itself.

### Visualization of model fits

Let 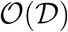 be a summary statistic of interest, that is an arbitrary function of a dataset 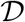 (e.g., the vestibular bias for a given bin of *s*_vis_ and visual reliability level, as per Figure 4 in the main text). For a given model, we generated the posterior predictive distribution of the group mean of 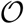 by following this bootstrap procedure:

• For *m* = 1,…, *M* = 100 iterations:

– Generate a synthetic group of *n* = 11 subjects by taking *n* samples from the individual posterior distributions of the model parameters.
– For each synthetic subject, generate a dataset 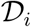 of simulated responses to the same trials experienced by the subject.
– Compute the group mean of the summary statistic across synthetic subjects, 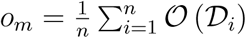.
• Compute mean and standard deviation of *o_m_*, which correspond to group mean and SEM of the summary statistic.

The shaded areas shown in the model fits figures in the main text are the posterior predictive distributions (mean ± SEM) of the summary statistics of interest.

### Model validation and recovery

We performed sanity checks and unit tests to verify the integrity of our code.

To test the implementation of our models, for a given observer (given model and parameter vector ***θ***) we tested the data simulation code (functions that simulate responses; used e.g. to generate figures) against the log likelihood code (functions that compute the log likelihood of the data). For a number of subjects and models we verified that, at the MLE solution, the log likelihood of the data approximated via simulation (by computing the probability of the responses via simple Monte Carlo) was ~ equal to the log likelihood of the data computed numerically. This ensured that our simulation code matched the log likelihood code, being a sanity check for both.

We performed a model recovery analysis to validate the correctness of our analysis pipeline, and assess our ability to distinguish models of interest using all tasks (‘joint fits’); see e.g. Acerbi, Ma, and Vijayakumar (2014); van den Berg et al. (2014). For computational tractability, we restricted our analysis to six observer models: the most likely four models for each different causal inference strategy (to verify our ability to distinguish between strategies), and, for the most likely model, its variants along the prior and noise factors (to verify whether we can distinguish models along those axes). Thus, we consider the following models: Fix-X-E, Bay-X-E, Bay/FFu-X-I, Fix/FFu-C-I, Fix-X-I, Fix-C-E (see main text for a description). We generated synthetic datasets from each of these six models, for all three tasks jointly, using the same sets of stimuli that were originally displayed to the 11 subjects. For each subject, we took four randomly chosen posterior parameter vectors obtained via MCMC sampling (as described in Section B), so as to ensure that the statistics of the simulated responses were similar to those of the subjects. Following this procedure, we generated 264 datasets in total (6 generating models × 11 subjects × 4 posterior samples). We then fit all 6 models to each synthetic dataset, yielding 1584 fitting problems. For computational tractability, we only performed maximum likelihood estimation (see Section B, with 50 restarts), as opposed to MCMC sampling, whose cost would be prohibitive for this number of fits. The analysis was otherwise exactly the same as that used for fitting the subject data. We then computed the fraction of times that a model was the ‘best fitting’ model for a given generating model, according to AICc (considering that AICc approximates LOO in the limit of large data).

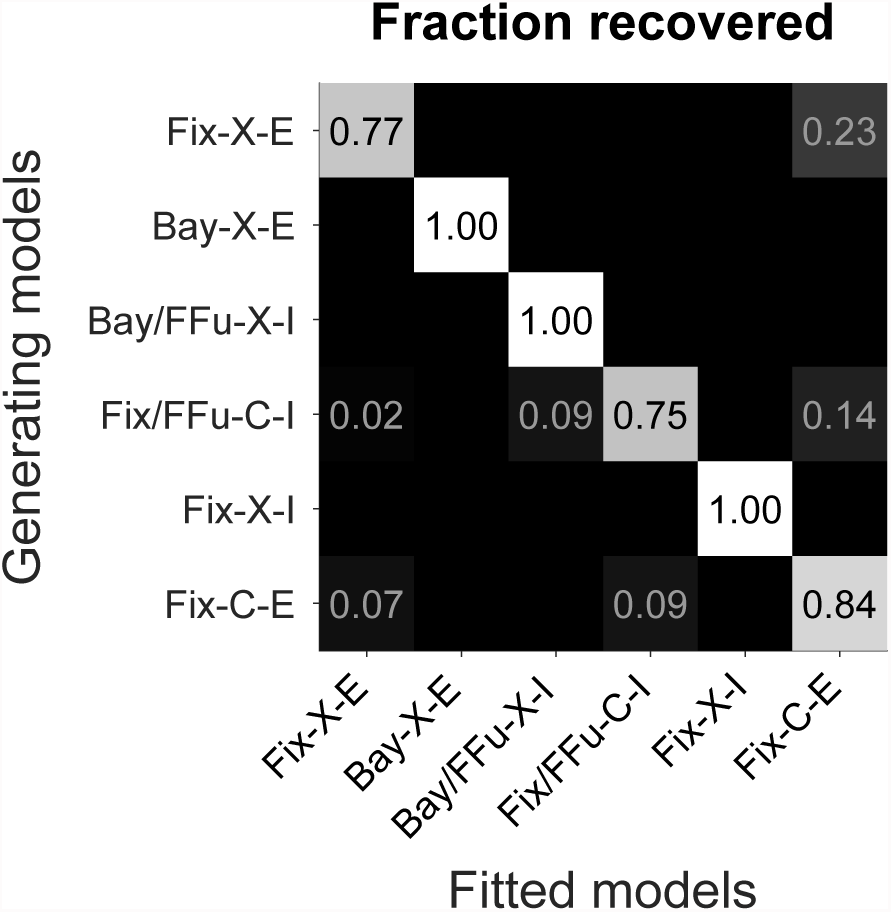

#### Model recovery analysis

Each square represents the fraction of datasets that were ‘best’ fitted from a model (columns), for a given generating model (rows), according to the AICc score. Bright shades of gray correspond to larger fractions. The bright diagonal indicates that the true generating model was, on average, the best-fitting model in all cases, leading to a successful model recovery.

We found that the true generating model was recovered correctly in 89.4% of the datasets on average (see above). This finding means that our models are distinguishable in a realistic setting, and at the same time validates the model fitting pipeline (as it would be unlikely to obtain a successful recovery in the presence of a substantial coding error). Since our model recovery method differs from the procedure used on subject data in the comparison metric (AICc via MLE, rather than LOO via MCMC), we verified on subject data that AICc and LOO scores were highly correlated across subjects (Adler & Ma, 2016). The Spearman’s rank correlation coefficient between the two metrics was larger than 0.99 for each of the sixteen models in the joint fits, providing strong evidence that results of our model recovery analysis would also transfer to the framework used for the subject data.

## Appendix C Absolute goodness of fit

In this appendix we describe a general method to compute absolute goodness of fit, largely based on the approach of Shen and Ma (2016).^4^

### Computing the absolute goodness of fit

Let ***X*** be a dataset of discrete categorical data grouped in *M* independent batches with *K* classes each, such that *X_jk_* is the number of observations for the *j*-th batch and the *k*-th class. We define *N_j_* = ∑*_k_ X_jk_* the number of observations for the *j*-th batch.

We assume that observations are ordered and independent, such that the distribution of observations in each batch *j* is the product of *Nj* categorical distributions with parameters ***p****_j_* = (*p_j_*_1_,…, *p_jK_*) (frequencies), such that the probability of the data is

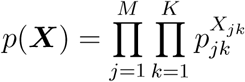

with unknown vectors of frequencies ***p****_j_.*

We assume that we have a model of interest *q* that predicts frequencies *q_jk_* for the observations, with ∑*_k_ q_jk_* = 1 for 1 ≤ *j* ≤ *M.* As a reference, we consider the chance model *q*^0^ with frequencies 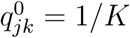.

We define the *absolute goodness of fit* of *q* as
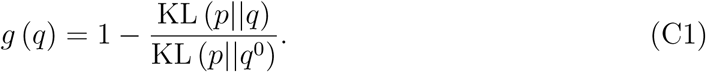

where KL (*p*||*q*) is the Kullback-Leibler divergence (also known as relative entropy) between a ‘true’ distribution *p* and an ‘approximating’ distribution *q.*

Importantly, *g*(*q*) = 0 when a model performs at chance, and *g*(*q*) ≤ 1, with *g*(*q*) = 1 only when the model matches the true distribution of the data. In other words, *g*(*q*) represents the fractional information gain over chance. Note that *g*(*q*) can be negative, in the unfortunate case that a model performs worse than chance.

As another important reference, we recommend to also compute the absolute goodness of fit 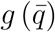 of the *histogram model* 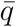, with frequencies defined from the empirical frequencies across batches as 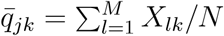, for 1 ≤ *j* ≤ *M* and *N* = ∑*_j_ N_j_*. A comparison between *g*(*q*) and 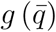 is informative of how better the current model is than a simple histogram of categorical observations collapsed across batches. In some circumstances, the chance model can be a straw model, whereas the histogram model may represent a more sensible reference point.

In order to estimate Eq. C1, we need to compute the relative entropy KL (*p||q*) between the data and a given distribution *q*,
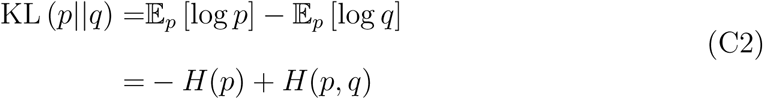

where the first term is the (negative) entropy of the data, and the second term is called the cross-entropy between *p* and *q.* We will show in the following sections that the negative cross-entropy is approximated by the cross-validated log likelihood of the data, LL_CV_ (*q*).

Combining Eq. C1 with our estimates of Eq. C2, we obtain
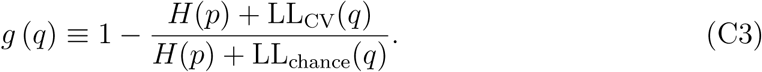

We show next how to estimate the entropy of the data, and prove that the negative cross-entropy between *p* and *q* is approximated by the cross-validated log likelihood.

### Entropy of the data

As noted in Shen and Ma (2016), the naïve plug-in estimator of the entropy of the data leads to a biased estimate of the entropy, and this bias can be substantial when the data are sparse (a few observations per batch). Instead, we use the Grassberger estimator of the entropy (Grassberger, 2003),
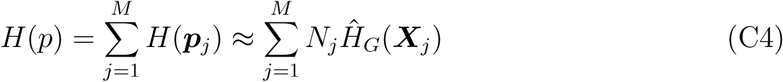

where the Grassberger estimator of the entropy per trial is defined as
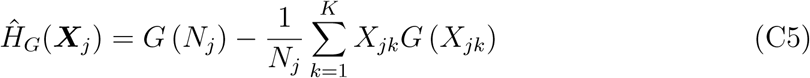

and *G*(*h*) for 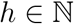 are Grassberger’s numbers defined as
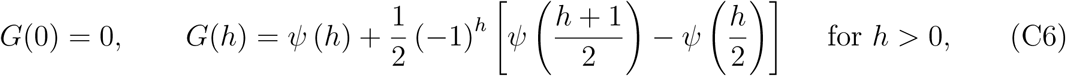

where *ψ* is the digamma function.

That is, our estimate of the negative entropy is
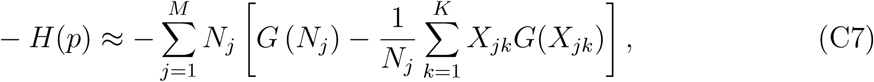

which is the same as Eq. 21 in Shen and Ma (2016), when restricted to the binomial case *(K* = 2), and after correcting for a typo (*N* in the denominator of their equation should read as *N_j_*).

### Cross-entropy

The estimated cross-entropy is
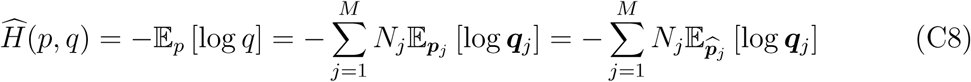

where in a slight abuse of notation we denoted with ***p_j_*** (resp., ***q_j_***) the categorical distributions associated to the data (resp., model) for the *j*-th batch. Crucially, since the expectations only involve *q*, 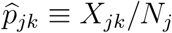 is an unbiased estimator of *p_jk_.*

Eq. C8 becomes
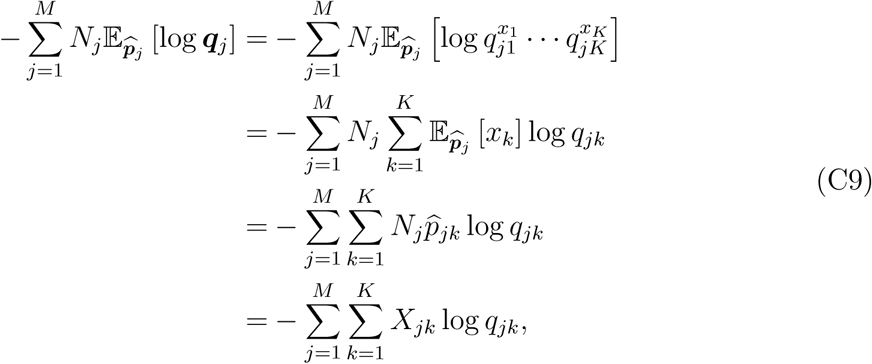

which is the negative log likelihood of the model, −LL(*q*).

Note that typically we also need to estimate the model parameters, and computing Eq. C9 on the same dataset used to estimate parameters will yield a biased estimate of the log likelihood (see e.g., Burnham & Anderson, 2003). Shen and Ma suggest to obtain an independent estimate of the log likelihood of the model via cross-validation, LL_CV_ (Shen & Ma, 2016). According to their method, model parameters are estimated on half of the data, and the log likelihood of the model (and also the entropy of the data) is evaluated with the other half of the data. As an improvement over their method, we advocate to estimate the expected log likelihood via leave-one-out (LOO) cross-validation score obtained via MCMC Vehtari et al. (2016). This will produce an unbiased estimator of the expected log likelihood, and allows to use all the available data to obtain a more robust estimate of the relative entropy.

In conclusion, our estimate for the cross-entropy is
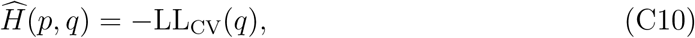

with LL_cv_(*q*) computed as the LOO score of the model, and it corresponds to Eq. 19 in Shen and Ma (2016).

## Appendix D LOO scores for all models

In this appendix we report tables of LOO scores for all models and subjects, which were used to perform group Bayesian Model Selection (BMS), the model comparison technique adopted in the main text. For each subject, LOO scores are shown relative to the LOO of the model with highest mean LOO across subject, which is printed in boldface. Models are ranked according to average LOO.

Summing (equivalently, averaging) LOO scores across subjects is a simple ‘fixed-effect’ model comparison analysis, in which all subjects are believed to belong to the same model. Results of the fixed-effect analysis differ in details from the group BMS, but the overall qualitative findings are analogous.

### Unity judgment task

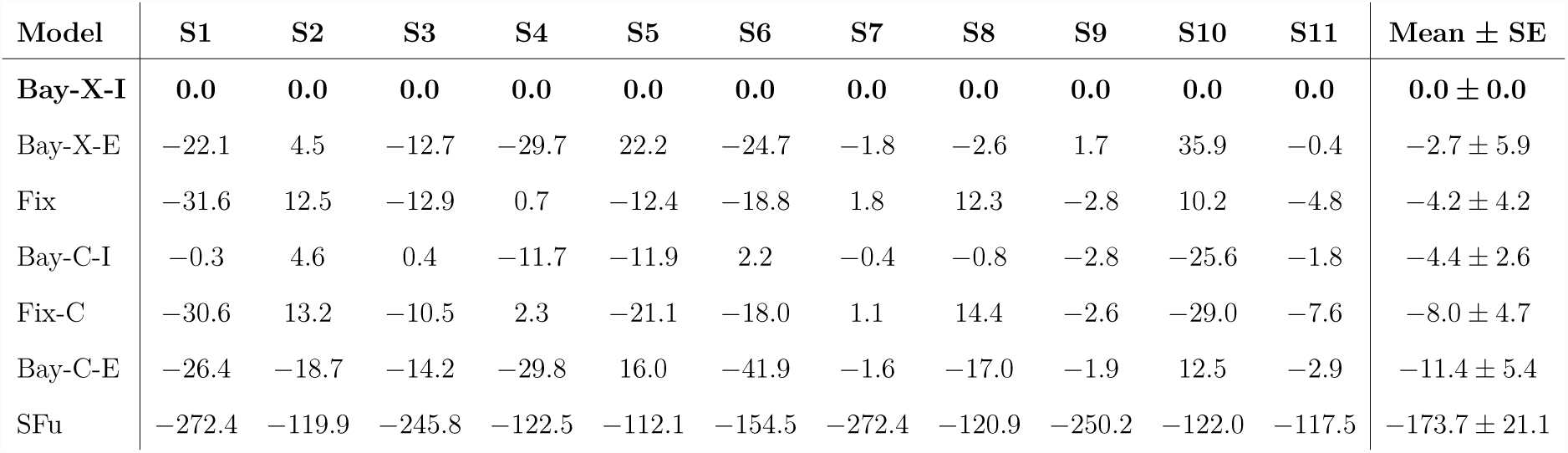

### Bimodal inertial discrimination task

| Model | S1 | S2 | S3 | S4 | S5 | S6 | S7 | S8 | S9 | S10 | S11 | Mean ± SE |
| --- | --- | --- | --- | --- | --- | --- | --- | --- | --- | --- | --- | --- |
| **Bay-X-E** | **0.0** | **0.0** | **0.0** | **0.0** | **0.0** | **0.0** | **0.0** | **0.0** | **0.0** | **0.0** | **0.0** | **0.0 ± 0.0** |
| Fix-X-E | -0.9 | 0.5 | -2.2 | -13.0 | -0.8 | 0.8 | -0.6 | 0.4 | 0.3 | 3.4 | 1.5 | -1.0 ± 1.3 |
| FFu-X-I | -0.7 | 0.9 | -3.5 | -11.3 | 0.4 | 1.3 | 1.2 | 1.6 | 1.5 | -12.3 | 1.1 | -1.8 ± 1.6 |
| Fix-X-I | -0.9 | 2.0 | -3.2 | -11.5 | 0.6 | 1.3 | 0.6 | 0.0 | 0.7 | -12.5 | 1.1 | -2.0 ± 1.6 |
| FFu-X-E | -0.2 | 0.9 | -3.6 | -10.2 | 0.6 | 1.5 | 0.9 | 1.5 | 1.4 | -18.8 | 1.2 | -2.3 ± 2.0 |
| Fix-C-E | -9.8 | -3.7 | 0.1 | -18.7 | -0.9 | -2.5 | -7.1 | 1.1 | -2.3 | 3.9 | 0.4 | -3.6 ± 1.9 |
| Bay-C-E | -10.5 | 0.3 | -0.6 | -5.7 | 0.6 | -1.9 | -11.8 | 0.1 | -1.8 | -5.4 | -3.2 | -3.6 ± 1.3 |
| Bay-X-I | -3.1 | -2.6 | -5.7 | -13.0 | -1.6 | -1.5 | 0.2 | -0.8 | 0.2 | -15.5 | 1.4 | -3.8 ± 1.7 |
| FFu-C-E | -20.1 | -22.1 | -9.9 | -34.7 | -14.8 | -21.9 | -31.9 | -6.0 | -2.4 | -57.7 | -2.1 | -20.3 ± 5.0 |
| FFu-C-I | -20.2 | -22.1 | -9.9 | -34.8 | -14.8 | -21.8 | -31.9 | -6.0 | -2.6 | -57.7 | -2.2 | -20.3 ± 5.0 |
| Fix-C-I | -20.2 | -22.1 | -9.9 | -34.8 | -14.8 | -21.9 | -30.6 | -6.8 | -3.1 | -57.8 | -2.3 | -20.4 ± 4.9 |
| Bay-C-I | -19.6 | -21.6 | -10.4 | -34.7 | -15.9 | -22.7 | -32.3 | -6.2 | -2.8 | -58.2 | -2.8 | -20.6 ± 5.0 |

### Joint fits

| Model | S1 | S2 | S3 | S4 | S5 | S6 | S7 | S8 | S9 | S10 | S11 | Mean ± SE |
| --- | --- | --- | --- | --- | --- | --- | --- | --- | --- | --- | --- | --- |
| **Fix-X-E** | **0.0** | **0.0** | **0.0** | **0.0** | **0.0** | **0.0** | **0.0** | **0.0** | **0.0** | **0.0** | **0.0** | **0.0 ± 0.0** |
| Fix-X-I | 7.3 | -7.2 | -4.4 | 31.9 | -16.1 | 23.7 | -40.6 | -26.6 | -38.6 | -20.4 | 2.4 | -8.0 ± 7.1 |
| Fix/FFu-X-E | -14.8 | -19.7 | -15.4 | 4.8 | -3.4 | 0.5 | -42.6 | -4.3 | -14.8 | -8.9 | 2.4 | -10.6 ± 4.0 |
| Fix-C-E | 0.4 | -9.2 | -1.9 | -14.3 | -41.9 | -7.9 | -3.5 | 0.9 | -6.5 | -46.7 | -10.7 | -12.8 ± 4.9 |
| Fix/FFu-X-I | -26.6 | -19.7 | -22.5 | 4.8 | -2.4 | 0.5 | -52.4 | -4.2 | -59.7 | -2.9 | 2.3 | -16.6 ± 6.7 |
| Bay-X-E | 17.1 | -34.2 | -6.2 | -31.3 | -25.8 | -20.6 | -9.5 | -128.6 | 12.7 | 12.3 | -6.2 | -20.0 ± 12.1 |
| Bay/FFu-X-E | -20.8 | -39.0 | -25.3 | 0.5 | 10.8 | -14.9 | -42.9 | -127.0 | -40.8 | 3.7 | 1.2 | -26.8 ± 11.6 |
| Fix/FFu-C-E | -14.2 | -21.0 | -17.9 | -20.5 | -47.6 | -6.0 | -44.5 | -3.3 | -19.0 | -103.6 | -8.5 | -27.8 ± 8.7 |
| Fix-C-I | -3.6 | -32.0 | -15.8 | -14.1 | -59.5 | 2.1 | -85.5 | -25.2 | -59.0 | -94.9 | -9.4 | -36.1 ± 10.1 |
| Fix/FFu-C-I | -25.6 | -21.1 | -22.0 | -20.6 | -47.7 | -6.1 | -86.0 | -3.4 | -59.1 | -103.7 | -8.6 | -36.7 ± 10.1 |
| Bay-C-E | 2.7 | -73.1 | -29.5 | -44.2 | -33.6 | -74.9 | -16.1 | -191.4 | -6.7 | -12.3 | -26.4 | -45.9 ± 16.4 |
| Bay/FFu-C-E | -36.4 | -77.3 | -47.1 | -31.0 | -15.4 | -45.3 | -90.7 | -206.7 | -72.9 | -74.3 | -11.8 | -64.4 ± 16.2 |
| Bay-X-I | -356.3 | -128.2 | -193.6 | -204.0 | -91.3 | -35.6 | -177.3 | -235.6 | -298.7 | -105.2 | -6.3 | -166.6 ± 32.2 |
| Bay-C-I | -462.0 | -222.2 | -318.1 | -231.3 | -158.8 | -77.6 | -319.6 | -338.0 | -488.1 | -259.2 | -51.2 | -266.0 ± 42.1 |
| Bay/FFu-X-I | -872.8 | -416.4 | -544.9 | -589.5 | -304.5 | -424.8 | -555.4 | -397.0 | -593.0 | -272.0 | -53.7 | -456.7 ± 64.2 |
| Bay/FFu-C-I | -888.7 | -445.3 | -556.3 | -611.2 | -340.1 | -441.8 | -551.3 | -396.2 | -625.7 | -351.2 | -69.8 | -479.8 ± 62.6 |

## Appendix E Model fits of full data for explicit inference task

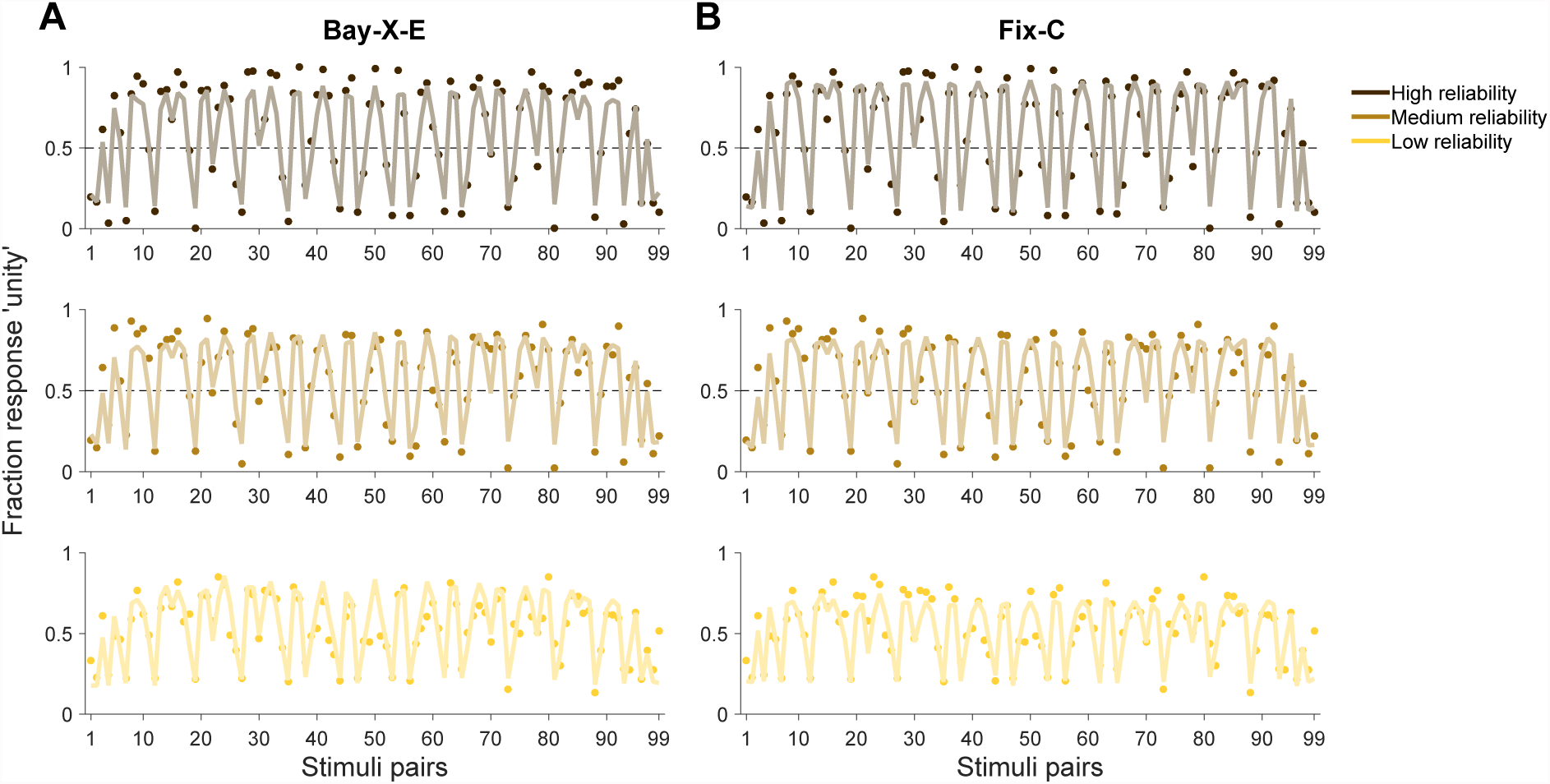

### Explicit inference task, full data

Results of the explicit inference (unity judgment) task, for two models of interest. Proportion of ‘unity’ responses for a given (*s*_vis_*, s*_vest_) heading direction pair (indexed from 1 to 99), and for different levels of visual cue reliability. Points are data, lines are model fits (average fit across subjects). Error bars are omitted for clarity. **A:** Best Bayesian model (Bay-X-E). **B:** Best fixed-criterion model (Fix-C). Neither model appears clearly superior across all noise levels (see main text).

## Footnotes

1 URL: <u>https://github.com/lacerbi/eissample</u>.

2 URL: <u>https://github.com/lacerbi/vbgmm</u>.

3 URL: <u>http://becs.aalto.fi/en/research/bayes/mcmcdiag/</u>.

3 URL: <u>https://github.com/lacerbi/gofit.</u>

